# Self-organization of kinetochore-fibers in human mitotic spindles

**DOI:** 10.1101/2021.11.11.468239

**Authors:** William Conway, Robert Kiewisz, Gunar Fabig, Colm P. Kelleher, Hai-Yin Wu, Maya Anjur-Dietrich, Thomas Müller-Reichert, Daniel Needleman

## Abstract

During eukaryotic cell division, chromosomes are linked to microtubules (MTs) in the spindle by a macromolecular complex called the kinetochore. The bound kinetochore microtubules (KMTs) are crucial to ensuring accurate chromosome segregation. Recent electron tomography reconstructions (Kiewisz et al. 2021) captured the positions and configurations of every MT in human mitotic spindles, revealing that many KMTs in these spindles do not reach the pole. Here, we investigate the processes that give rise to this distribution of KMTs using a combination of analysis of the electron tomography reconstructions, photoconversion experiments, quantitative polarized light microscopy, and biophysical modeling. Our results indicate that in metaphase, KMTs grow away from the kinetochores along well-defined trajectories, continually decreasing in speed as they approach the poles. The locations of KMT minus ends, and the turnover and movements of tubulin in KMTs, are consistent with models in which KMTs predominately nucleate de novo at kinetochores and are inconsistent with substantial numbers of non-KMTs being recruited to the kinetochore in metaphase. Taken together, this work leads to a mathematical model of the self-organization of kinetochore-fibers in human mitotic spindles.

## INTRODUCTION

When eukaryotic cells divide, a spindle composed of microtubules (MTs) and associated proteins assembles and segregates the chromosomes to the daughter cells (Strasberger et al 1880, McIntosh 2012, Heald and Khodjakov 2015, Petry et al. 2016, Prosser and Pelletier 2017, Oriola et a. 2018, Anjur-Dietrich et al. 2021). A macromolecular protein complex called the kinetochore binds each sister chromatid to MTs in the spindle thereby bi-orienting the two sisters to ensure they segregate to opposite daughter cells (McDonald et al. 1992, McEwen et al. 1997, Yoo et al. 2017, Monda et al. 2018 Rieder 1982, Maiato et al. 2004b, Mussachio et al, 2017, Pesenti et al. 2018, Monda and Cheeseman 2018 DeLuca et al. 2011, Redemann et al. 2017, Long et al. 2019). An MT whose plus end is embedded in the kinetochore is referred to as a kinetochore microtubule (KMT) and the collection of all KMTs associated with an individual kinetochore is called a kinetochore-fiber (K-Fiber). The kinetochore-microtubule interaction stabilizes KMTs and generates tension across the sister chromatid pair (Brinkley and Cartwright 1975, Gorbsky and Borisy 1989, Nicklas and Ward 1996, DeLuca et al. 2006, Cheeseman et al. 2006, Tanaka and Desai 2008, Akiyoshi et al 2010, Kabeche and Compton 2013, Cheerambathur et al. 2017, Monda and Cheeseman 2018, Steblyanko et al. 2020 Warren et al. 2020). Modulation of the kinetochore-microtubule interaction is thought to be important in correcting mitotic errors (DeLuca et al. 2011, Godek et al. 2015, Funabiki 2019, Long et al 2019). Kinetochore-microtubule binding is thus central to normal mitotic progression and correctly segregating sister chromatids to opposite daughter cells (Cimmini et al. 2001, Chiang et al. 2010, Auckland and McAinsh 2015, Lampson and Grishchuk 2017, Dudka et al. 2018). Chromosome segregation errors are implicated in a host of diseases ranging from cancer to development disorders such as Downs’ and Turners’ Syndromes (Touati and Wassmann 2016, Compton 2017, Jo et al 2021).

The lifecycle of a KMT consists of its recruitment to the kinetochore, its subsequent motion, polymerization and depolymerization, and its eventual detachment from the kinetochore. The initial recruitment of an MT to the kinetochore can either occur by a non-KMT being captured by the kinetochore, or by de-novo nucleation of a KMT at the kinetochore (Telzer et al. 1975, Mitchinson and Kirschner 1985a, Mitchinson and Kirschner 1985b, Huitorel and Kirschner 1988, Heald and Khodjakov 2015, LaFountain and Oldenborug 2014, Petry 2016, Sikirzhyski et al. 2018, David et al. 2019, Renda and Khodjakov 2021). The relative importance of these two pathways throughout mitosis in human cells remains unknown. The plus-ends of KMTs can polymerize and depolymerize while remaining attached to the kinetochore, leading to a net flux of tubulin through the K-Fiber from the kinetochore towards the spindle pole (Rieder and Alexander 1990, Mitchinson and Salmon 1992, Zhai et al. 1995, Waters et al. 1996, Khodjakov et al. 2003, Gabbe and Heald 2004, McIntosh et al. 2012, Steblyanko et al. 2020, DeLuca et al. 2011, Elting et al. 2014, Elting et al. 2017, Neahring et al. 2021, Risteski et al. 2021). For human cells in metaphase, it is unclear to what extent these motions are due to movement of entire K-Fibers, movement of individual KMTs within a K-Fiber, or movement of tubulin through individual KMTs. Finally, when KMTs detach from the kinetochore, they become non-KMTs by definition. The regulation of KMT detachments is thought to be important for correcting improper attachments and ensuring accurate chromosome segregation (Tanaka et al. 2002, Bakhoum et al. 2009, DeLuca et al. 2011, Godek et al. 2015, Krenn and Mussachio 2015, Lampson and Grishuk 2017, Funabiki 2019, Long et al 2019,). KMT detachments typically occur with a time scale of ∼5 mins in metaphase in human mitotic cells (Kabeche and Compton 2013). How these processes – KMT recruitment, motion, polymerization and depolymerization, and detachment – lead to the self-organization of K-Fibers remains incompletely understood.

In a companion paper, we used serial-section electron tomography to reconstruct the locations, lengths, and configurations of MTs in metaphase spindles in HeLa cells (Kiewisz et al 2021). These whole spindle reconstructions can unambiguously identify which MTs are bound to the kinetochore and measure their lengths, providing a remarkable new tool for the study of KMTs. Strikingly, many KMTs do not reach all the way to the pole. Here, we sought to combine the electron tomography spindle reconstructions with live-cell experiments and biophysical modeling to characterize the lifecycle of KMTs in metaphase spindles in HeLa cells. The electron tomography reconstructions revealed that only ∼50% of KMTs have their minus ends at spindle poles. We used photoconversion experiments to measure the dynamics of KMTs, which revealed that while their stability does not spatially vary, their speed is greatest in the middle of the spindle and continually decreases closer to poles. We next show that the orientations of MTs throughout the spindle, measured by electron tomography and polarized light microscopy, can be quantitively explained by an active liquid crystal theory in which the mutual interactions between MTs cause them to locally align with each other. This argues that KMTs tend to move along well-defined trajectories in the spindle. We show that the distribution of KMT minus ends along these trajectories (measured by electron tomography) is only consistent with the motion and turnover of KMTs (measured by photoconversion) if KMTs predominately nucleate at kinetochores. Taken together, these results lead us to construct a model in which KMTs nucleate at the kinetochore, grow and slow down as they move along their trajectories toward poles, undergo minus end depolymerization near the pole and detach from the kinetochore at a constant rate. Such a model of K-Fiber self-organization can quantitively explain the lengths, locations, configurations, motions, and turnover of KMTs throughout metaphase spindles in HeLa cells.

## RESULTS

### Many KMT minus ends are not at the pole

We first analyzed a recent cellular tomography electron microscopy (EM) reconstruction data set which captured the trajectories of every MT in the mitotic spindle of three HeLa cells (Kiewisz et al., 2021). We defined KMTs as MTs with one end near a kinetochore in the reconstructions and assigned the plus end to the end at the kinetochore and the minus end to the opposite end of the MT (Figure 1A). KMT minus ends are located throughout the spindle, with approximately 51% of them more than 1.5μm away from the pole (Figure 1B). KMT minus ends are distributed throughout individual K-Fibers (Figure 1C), indicating that the processes that lead to a broad distribution of KMT minus end locations can occur at the level of individual kinetochores. We wanted to know how the observed distribution of KMT minus end locations results from the behaviors of KMTs. This requires understanding the life cycle of a metaphase KMT, namely (Figure 1D):

1. How are KMTs recruited to kinetochores? To what extent are they nucleated de novo at the kinetochore vs. resulting from non-KMTs being captured from the bulk of the spindle?
2. How do KMTs move and grow? What are their growth trajectories and the minus end speeds?
3. How do KMTs detach from kinetochores?

**Figure 1:**
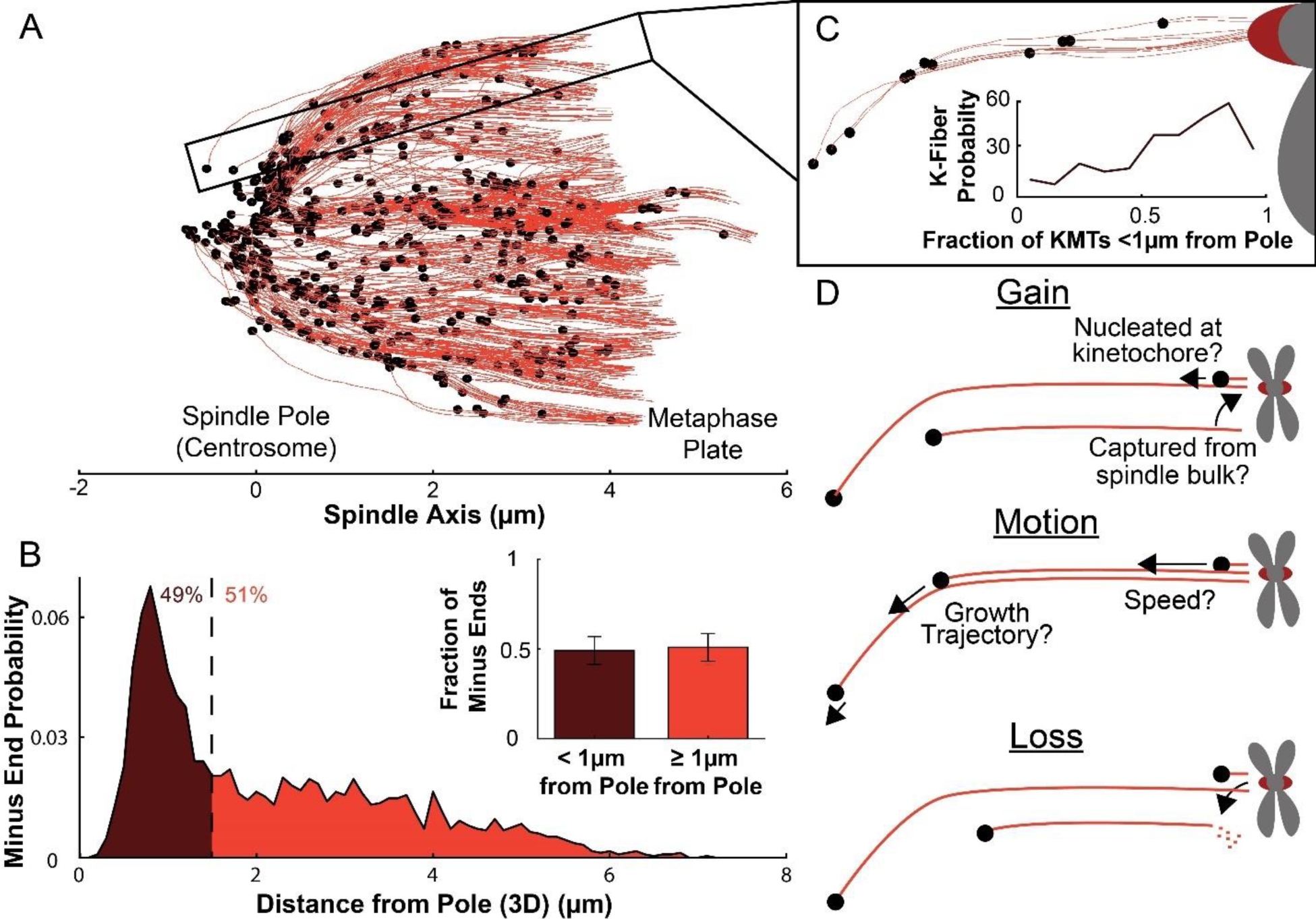
Many KMT minus ends are not in the vicinity of the pole. A) A sample half spindle showing the KMTs from the EM ultrastructure. KMTs are shown in red while minus ends are shown in black. The spindle pole lies at 0µm on the spindle axis while the metaphase plate is between 4-6 µm on the spindle axis. B) The frequency of 3D minus end distance from the pole. Inset: the fraction of minus ends within 1µm of the pole. C) A sample k-Fiber. Again, KMTs are shown in red, minus ends are shown in black. The large red circle is the kinetochore. Inset: probability of k-Fiber with fraction of KMTs near the pole D) Schematic representation of models of KMT gain, motion and loss.

We sought to answer these questions with a series of live-cell experiments, further analysis of the spindle reconstructions obtained from electron tomography, and mathematical modeling.

### The fraction of slow-turnover tubulin measured by photoactivation matches the fraction of tubulin in KMTs measured by electron tomography

To understand how the motion and turnover of KMTs results in the observed ultrastructure, we first sought to characterize the motion and stability of KMTs throughout the spindle. To that end, we constructed a Hela line stably expressing CENP-A:GFP to mark kinetochores and mEOS3.2:alpha tubulin to mark MTs. After photoconverting a line of tubulin in the spindle, the converted tubulin moves poleward and fades over time (Figure 2A) (Mitchinson 1989, DeLuca 2010, Kabeche and Compton 2013, Yu et al. 2019, Steblyanko et al. 2020).

**Figure 2:**
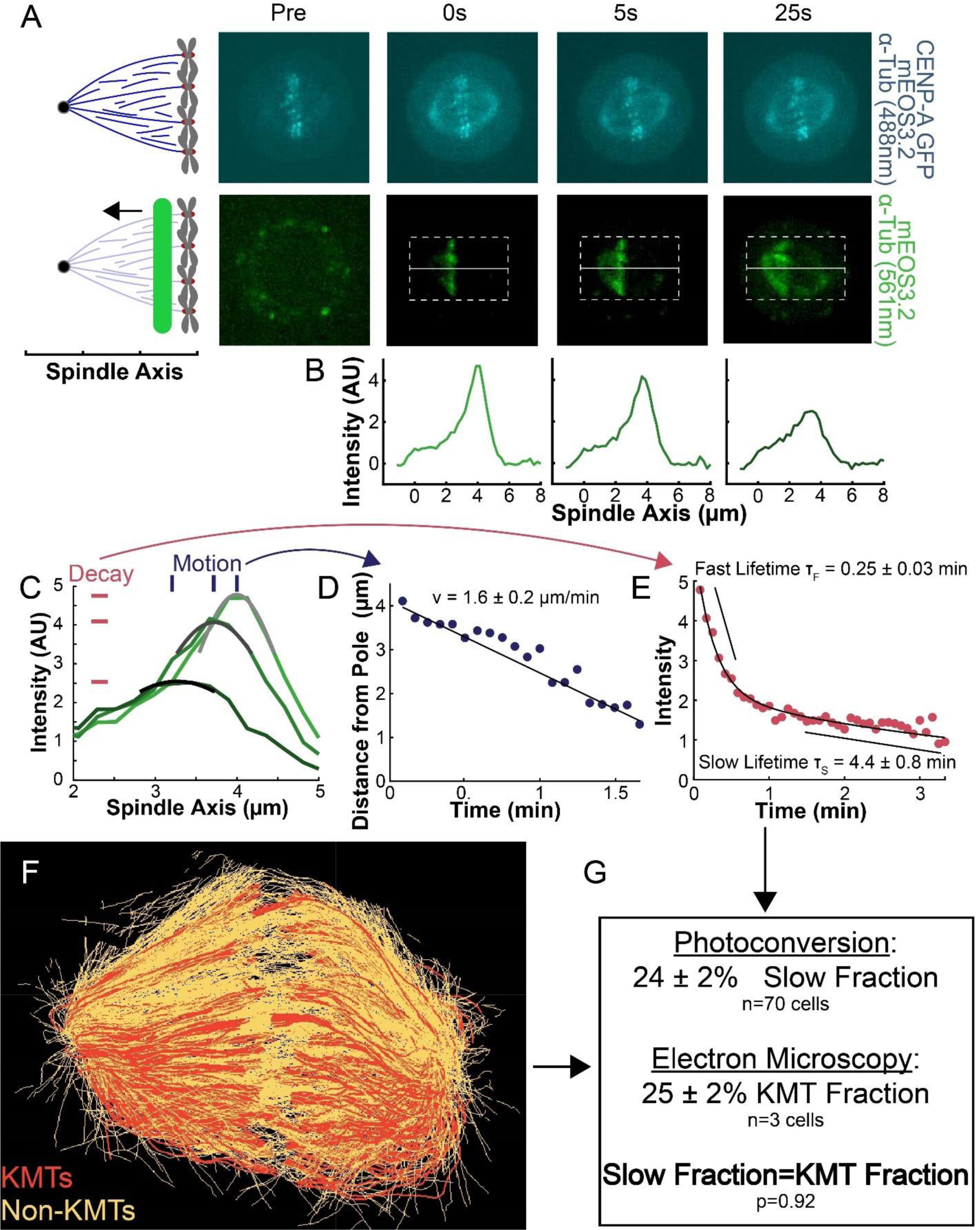
Photoconversion of spindle tubulin in live HeLa cells. A) Pre-converted frame showing CENPA-GFP and mEOS3.2-alpha tubulin. 488nm, 300ms exposure, 5s frame rate. Time stamps show pre photoconversion, 0s, 5s and 25s after photoconversion. Post-conversion frame showing mEOS3.2-alpha tubulin after exposure to 40nm light. 561nm, 500ms exposure, 5s frame rate. B) Line profile pulled from the dotted box shown in B. The intensity is corrected for background from the opposite side of the spindle (see methods) C) Line profiles (shades of green) fit to Gaussian profiles (shades of grey) at 0s, 5s and 25s. Lighter shades are earlier times. The solid line on the fit represents the fit pixels D) Blue dots: fit position of the line profile peak from the sample cell shown in A, B, and C over time. Black line: linear fit to the central position of the fit peak over time. E) Red dots: fit height of the line profile peak from the sample cell shown in A, B, and C over time. Black line: dual-exponential fit to the fit height of the peak over time. F) Sample ultrastructure from a single EM spindle (Kiewisz et al., 2021). KMTs are shown in red, non-KMTs yellow). G) Comparison between the mean slow fraction from the photoconversion data (24±2%, n=70 cells) and the fraction of KMTs(25±2%, n=3 cells) from the EM data. The two means are statistically indistinguishable with p=0.92 on a Student’s t-test.

To measure the speed and turnover of MTs, we first projected the intensity of the photoconverted tubulin onto the spindle axis (Figure 2B) (Kabeche and Compton 2013). We then fit the resulting peak to a Gaussian to track the motion of its center position and decay of its height over time (Figure 2C). We fit the position of the peak center over time to a line to determine the speed of tubulin movement in the spindle (Figure 2D). We then corrected the peak heights for bleaching by dividing by a bleaching reference (Figure 2s1) and fit the resulting time course to a dual-exponential decay to measure the tubulin turnover dynamics (Figure 2E) (DeLuca 2010).

Since the tubulin turnover is well-fit by a dual-exponential decay, it suggests that there are two subpopulations of MTs with different stabilities in the spindle, as previously argued for many model systems (Brinkley 1975, Salmon et al. 1976, Lambert and Bajer 1977, Rieder and Bajer 1977, Rieder 1981, Cassimeris et al. 1990, DeLuca et al. 2010). In prior studies, the slow-turnover subpopulation has typically been ascribed to the KMTs, while the fast-turnover subpopulation has typically been ascribed to the non-KMTs (Zhai et al. 1995, DeLuca 2010, Kabeche and Compton 2013). However, it is hypothetically possible that a portion of non-KMTs are also stabilized, due to bundling or some other mechanism (Tipton et al. 2021). To gain insight into this issue, we generated a cell line with SNAP-centrin to mark the poles and mEOS3.2:alpha tubulin to mark MTs and performed photoconversion experiments on a total of 70 spindles. We compared the fraction of tubulin in KMTs, 25±2% (n = 3), measured by electron tomography (in which a KMT is defined morphologically as a MT with one end embedded in a kinetochore; Figure 2F; Kiewisz, et al., 2021) to the fraction of the slow-turnover subpopulation measured from photoconversion experiments, 24±2% (n = 70). Since these two fractions are statistically indistinguishable (Figure 2G, p=0.92 on a Students’ t-test), we conclude that the slow-turnover subpopulation are indeed KMTs, and that there is not a significant number of stabilized non-KMTs.

### KMT speed is spatially varying while both KMT and non-KMT stability are uniform in the spindle bulk

We next explored the extent to which the speed and stability of MTs changed throughout the spindle (Burbank et al. 2007, Yang et al. 2008). To do this, we compared photoconversion results from lines drawn at different position along the spindle axis. After photoconverting close to the center of the spindle (∼4.5µm from the pole), the resulting line of marked tubulin migrated towards the pole (Figure 3A). This poleward motion was less evident when we photoconverted a line halfway between the kinetochores and the pole (Figure 3B), and barely visible when we photoconverted a line near the pole itself (Figure 3C). Tracking the subsequent motions of these photoconverted lines in different regions revealed clear differences in their speeds (Figure 3D), while their turnover appeared to be similar (Figure 3E). To quantitively study this phenomenon, we photoconverted lines in 74 different spindles, at various distances from the pole and measured the speed and turnover times at each location. Combining data from these different spindles revealed that average speed of the photoconverted lines increased with increasing distance from the pole (Figure 3F; Slope=0.20±0.07(µm/min)/µm, p=0.004), while both the KMT (Figure 3G; Slope=-0.03±0.05(1/min)/µm, p=0.23) and non-KMT (Figure 3H; Slope=0.0±0.2 (1/min)/µm, p=0.44) turnover were independent of distance from the pole. These results suggest that the speed of the KMTs is faster the further they are from the pole, and that the stability of KMTs and non-KMTs are constant throughout the spindle.

**Figure 3:**
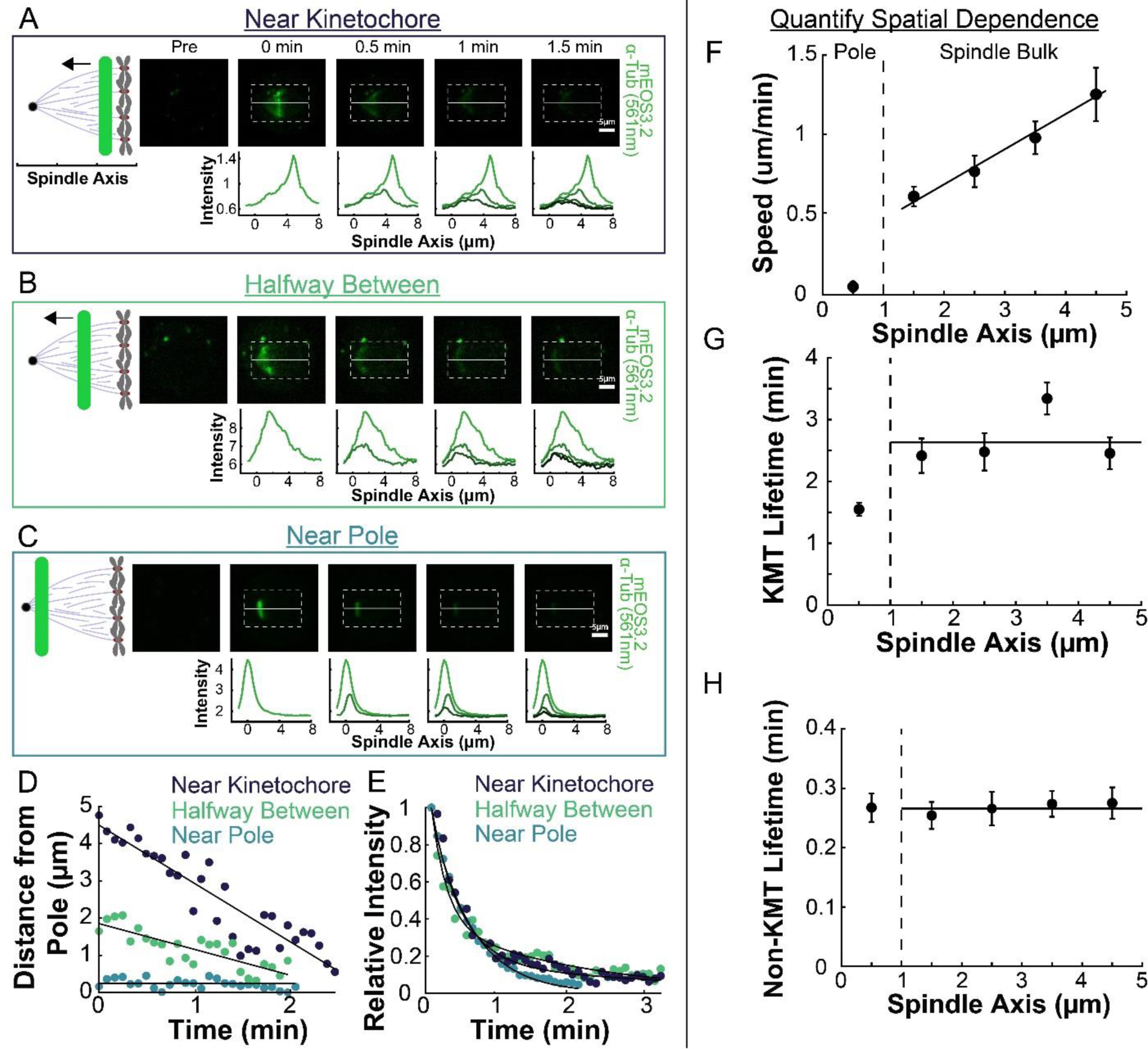
Spatial dependence of photoconversion parameters. A) Sample photoconverted frames (561nm, 500ms exposure, 5s frame rate) and line profiles from a line drawn near the kinetochore. B) Sample photoconverted frames and line profiles from a line drawn halfway between the kinetochores and the pole. C) Sample photoconverted frames and line profile from a line drawn near the pole. D) Linear fits to the central position of the peaks from A, B and C to measure the line speed.s E) Dual-exponential fits to the intensity of the line in A, B and C to measure the KMT and non-KMT lifetimes. F) Line speed vs. initial position of the line drawn on the spindle axis. The area near the pole and in the spindle bulk are marked, divided by a dashed line at 1µm. Error bars are standard error of the mean. (0-1µm: n=18; 1-2µm: n=14; 2-3µm: n=15; 3-4µm: n=16; 4-5µm: n=6) G) KMT lifetime vs. initial position of the line drawn on the spindle axis. H) Non-KMT lifetime vs. initial position of the line drawn on the spindle axis.

### KMTs and non-KMTs are well aligned in the spindle

To connect the static ultrastructure of KMTs (visualized by electron tomography) to the spatially varying KMT speeds (measured by photoconversion), we next sought to better characterize the orientation and alignment of MTs in the spindle. We started by separately analyzing the non-KMTs and KMTs (Figure 4A) in all three electron tomography reconstructions (Figs 4s1,4s2), and found that all MTs overwhelmingly lie on trajectories in the spindle axis-radial axis plane (Figure 4s3). We therefore projected all MTs into this plane and calculated the average orientation, 〈θ〉, in the spindle for both non-KMTs (Figure 4B) and KMTs (Figure 4C). The orientations of non-KMTs and KMTs were very similar to each other throughout the spindle, as can be seen by comparing the mean orientation of both sets of MTs along the spindle axis (Figure 4D). Thus, the non-KMTs and KMTs align along the same orientation field in the spindle.

**Figure 4:**
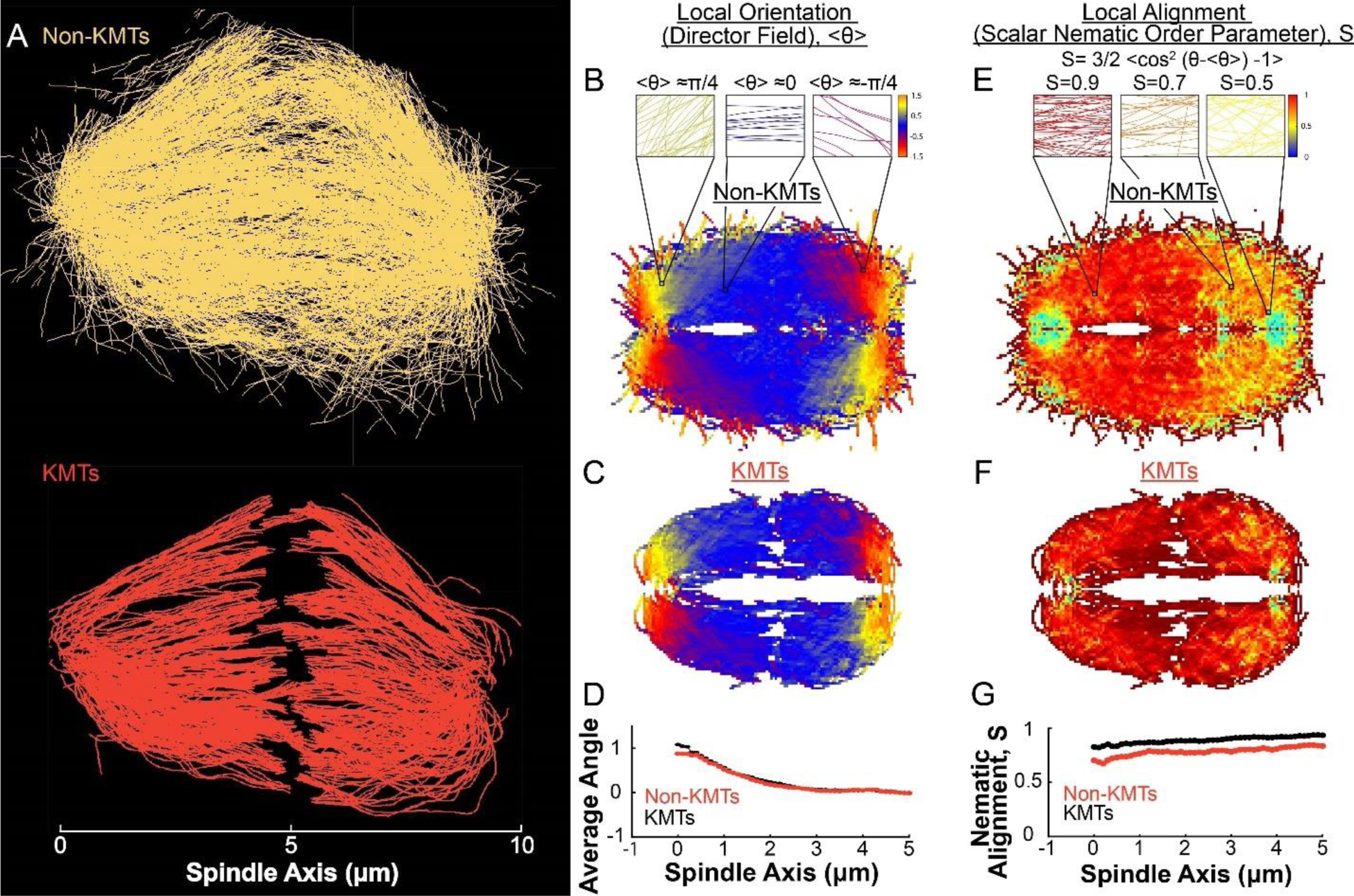
Measuring nematic alignment of non-KMTs and KMTs. A) Sample from a single EM reconstruction of non-KMTs (yellow) and KMTS (red). B) Mean local orientation of non-KMTs average over all theta along the spindle axis. Sample calculations of the local orientation in three representative pixels are shown above (yellow θ=π/4, blue θ=0m red θ=-π/4). C) Mean local orientation of KMTs average over all theta along the spindle axis. D) Averaged orientation angle of KMTs (red) and non-KMTs (black) along the spindle axis. E) Local alignment of the non-KMTs. Sample calculation of the local orientation in three representative pixels are shown above (yellow θ=π/4, blue θ=0m red θ=-π/4). F) Local alignment of the KMTs. G) Average alignment of the non-KMTs (black) and KMTs (red).

The above analysis addresses how the average orientation of MTs varies throughout the spindle. We next sought to quantify the degree to which MTs are well aligned along these average orientations. This is conveniently achieved by calculating the scalar nematic order parameter, *S* = 3⁄2 〈cos^2^(θ − 〈θ〉) − 1〉, which would be 1 for perfectly aligned MTs and 0 for randomly ordered MTs (de Gennes and Prost 1993). We calculated *S* for both non-KMTs (Figure 4E) and KMTs (Figure 4F) throughout the spindle. Both sets of MTs are well aligned throughout the spindle (Figure 4G) with 〈*S*〉 = 0.90 ± 0.01 for KMTs and 〈*S*〉 = 0.78 ± 0.01 for non-KMTs. The strong alignment of MTs in the spindle along the (spatially varying) average orientation field suggests that MTs in the spindle tend to move and grow along this orientation field.

We next calculated the orientation field of MTs in Hela spindles by averaging together data from both non-KMTs and KMTs from all three EM reconstructions by rescaling each spindle to have the same pole-pole distance and radial width (Figure 5A). We sought to test if the resulting orientation field was representative by obtaining data on additional Hela spindles. Performing significantly more large-scale EM reconstructions is prohibitively time consuming, so we turned to an alternative technique: the LC-Polscope, a form of polarized light microscopy that can quantitively measure the optical slow axis (i.e. the average MT orientation) with optical resolution (Oldenbourg et al. 1998) We averaged together live-cell LC-Polscope data from eleven Hela spindles and obtained an orientational field (Figure 5B) that looked remarkably similar to the one measured by EM (compare Figure 5A and 5B).

**Figure 5:**
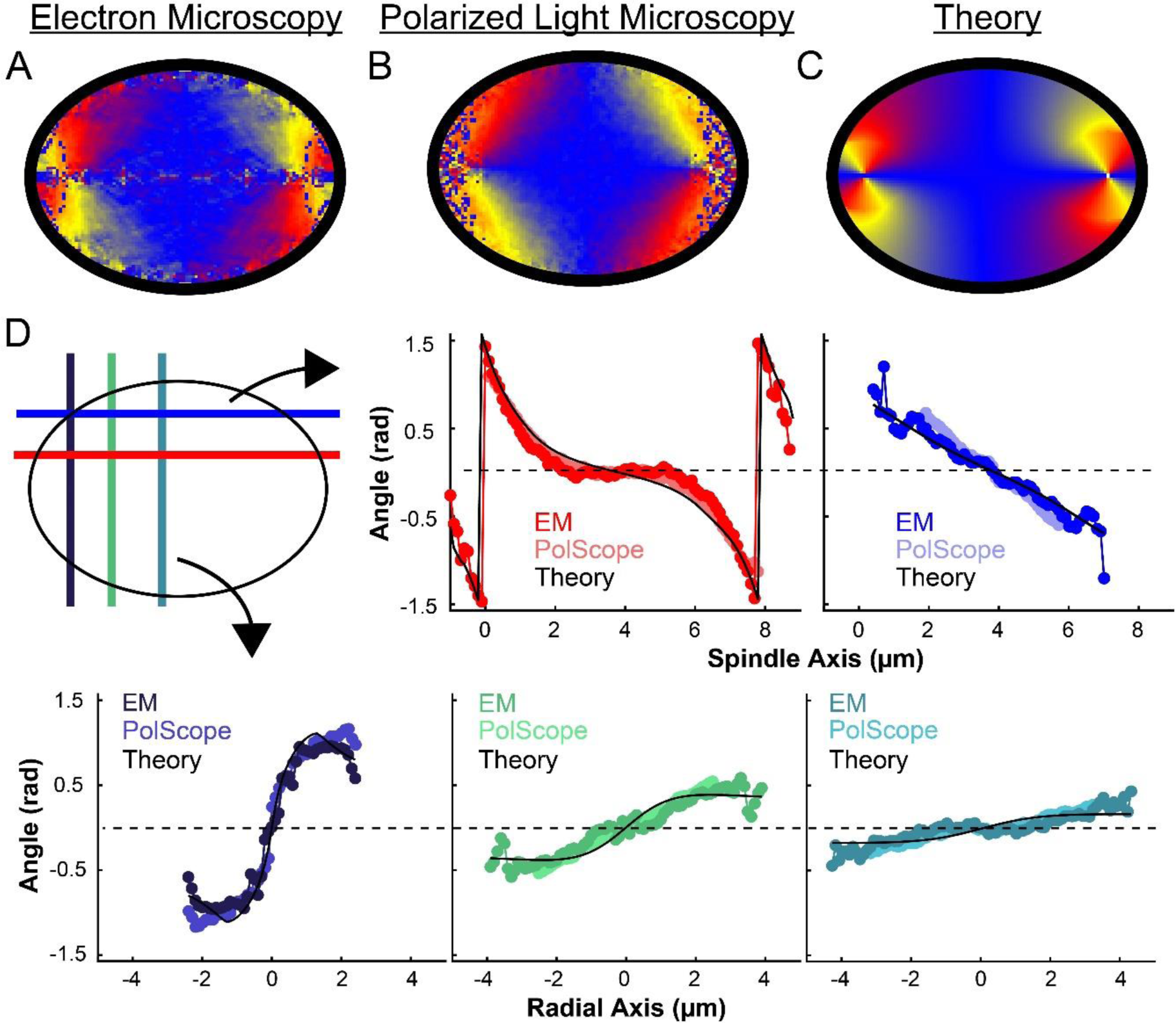
Experiment and theory of the orientation field of MTs in HeLa spindles. A) Orientation field of MTs from averaging electron microscopy (EM) reconstructions from three spindles. B) Orientation field of MTs from averaging polarized light microscopy (LC-PolScope) data from eleven spindles. C) A theoretical model of the spindle geometry with tangential anchoring at the elliptical spindle boundary and point defects at the poles. D) Average angle along narrow cuts parallel to the spindle and radial axis (red-lower spindle cut, blue-upper spindle cut purple-radial cut near pole, green-radial cut halfway between pole and kinetochore, teal-radial cut near kinetochore) shows close agreement between orientations from EM, polscope, and theory (black lines).

Previous work has shown that the internal dynamics and orientation of MTs in *Xenopus* egg extract spindles can be quantitively explained by an active liquid crystal theory (Brugués and Needleman 2014, Oriola et al. 2020). In this theory, the morphology of the spindle results from the local interactions of MTs with each other (mediated by molecular motors and other cross-linkers), which cause MTs to locally align relative to each other. A remarkable prediction of this theory is that the orientations of MTs in the spindle satisfy Laplace’s equation, ∇^2^θ = 0, where θ is the average local orientation of MTs. Thus, this theory predicts that the orientations of MTs throughout the spindle are completely determined by the spindle’s boundary and topological defects and, once those are specified, do not depend on parameters, such as those representing the MTs interactions or dynamics. We tested if this same framework can accurately describe Hela spindles by calculating the expected MT orientation field with tangential anchoring at the spindle boundary. In this calculation we adjusted the location and size of the two point defects, with a best fit placing them near the centrosomes as expected (Figure 5C). The theoretically predicted orientation field is remarkably similar to the orientation fields experimentally measured with EM and LC-Polscope (Figure 5D). Displacing the point defects to alternative locations, such as at the spindle periphery, results in substantially worse fits (Figure 5S1).

The agreement between the active liquid crystal theory, EM and LC-Polscope argues that the orientation of MTs in Hela spindles are determined by MTs locally interacting with each other. This, in turn, suggests that MTs in Hela spindles tend to grow and move along the direction set by the orientation field.

### The distribution of KMT minus ends along streamlines constrains models of KMT behaviors

We next explored in more detail the implication that KMTs grow and move along the orientation field of the spindle. If the trajectories of KMTs are confined to lie along the orientation field, then their minus ends will trace out paths on streamlines which lie tangent to the director field as they move towards the pole. We define a coordinate *s* as the distance from the pole along the streamlines, with *s* = 0 at the pole itself for all streamlines. We started by considering the locations of KMT minus ends on such streamlines. For each of the three individual reconstructed spindles, we fit the average MT orientations to the director field predicted by the active liquid crystal theory with two point defects and tangential anchoring along the spindle boundary (Figure 6s1). Then, for each KMT in each spindle, we integrated the fit director field from the KMT’s minus end to the associated spindle pole to find the streamline trajectory and calculated the corresponding location as the arc length along that streamline (Figure 6A). We combined data from the three electron tomography reconstructions to construct the density distribution along streamlines of KMT minus ends whose plus ends were upstream of that position (Figure 6B, see modeling supplement). This distribution peaks roughly 1µm away from the pole and is flat in the spindle bulk.

**Figure 6:**
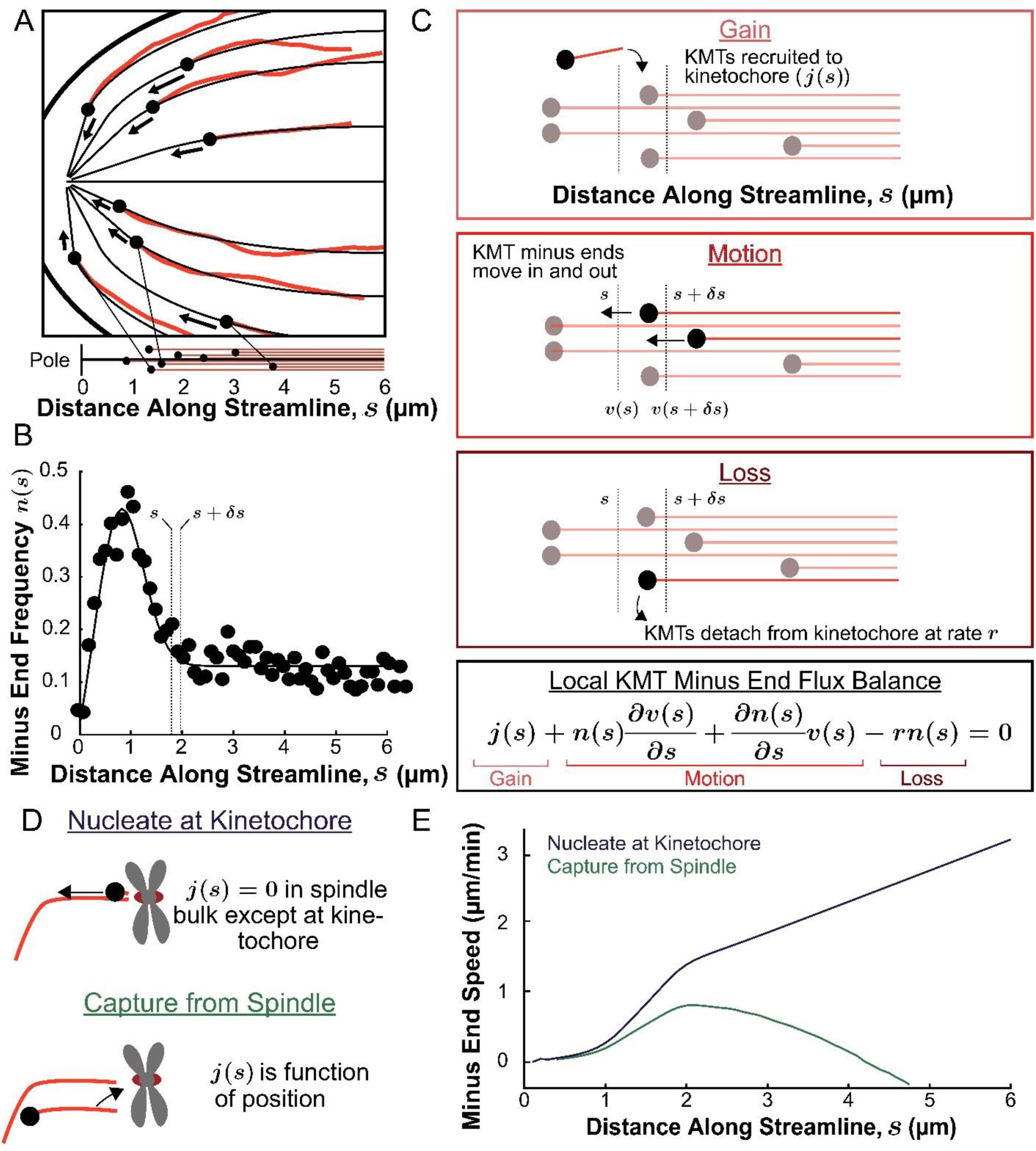
Predicting the KMT minus end speeds from the steady state distribution of minus ends along streamlines. A) Eight representative KMTs from the electron microscopy reconstruction (red), with their minus ends (black dots) and the streamlines (thin black lines) these minus ends are located on. The distance of these minus ends along the streamlines, *x*, are depicted (lower). B) Binned histogram, combining data from all three EM reconstructions, of the frequency along streamlines of KMT minus ends whose plus ends were upstream of that position. Histogram is fit to a Gaussian peaked near the pole and a constant in the spindle bulk (black line). C) Schematic depicting cartoon representations of KMT recruitment, minus end position and KMT detachment. The three cartoons depict KMT gain, (*j*(*s*)), KMT minus end motion in *n*(*s* + *δs*)*v*(*s* + *δs*)), KMT minus end motion out (*n*(*s*)*v*(*s*)) and MT loss (rk). Balancing these fluxes gives the mass conservation equation 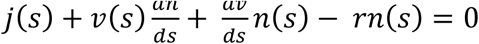 Cartoonshowing two models of KMT nucleation 1. nucleate at the kinetochore where *j*(*s*) = 0everywhere except at the kinetochore and 2. capture from spindle where *j*(*s*) is a function of position in the spindle. E) Comparison of the predictions of KMT minus end speeds in the nucleate at kinetochore and capture from spindle models.

The assumption that KMTs lie along streamlines suggests that this distribution of KMT minus ends results from the balance of three processes (Figure 6C): 1) If a non-KMT whose minus end is at position *s* along a streamline grows such that its plus end binds a kinetochore, then that non-KMT is recruited to become a KMT. This results in the addition of a new KMT minus end appearing at position *s*, which occurs with a rate *j*(*s*); 2) Microtubule minus ends move towards the pole with a speed, *v*(*s*), that may vary with position along the streamline; 3) When a KMT whose minus end is at position *s* along a streamline detaches from the kinetochore it becomes a non-KMT (by definition). This results in the loss of a KMT minus end at position *s*, which occurs at a rate *r.* The observation that the turnover rates of KMTs, as measured by photoactivation, is uniform throughout the bulk of the spindle (Figure 3G) argues that the detachment rate, *r* does not depend on the position along a streamline.

If the measured distribution of KMT minus ends (Figure 6B) is at steady-state, then the fluxes from the three processes described above – gain, movement, and loss – must balance at all locations along streamlines (Figure 6C), leading to:

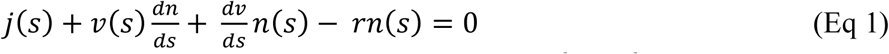

Where *n*(*s*), is the density of KMT minus ends at position *s,* and 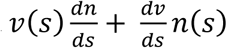 is the flux that results from the difference between KMT minus ends moving in and out of position *s*. Thus, Eq. 1 specifies a relationship between the distribution of KMT minus ends, *n*(*s*), the spatially varying speed of KMT minus ends, *v*(*s*), and rate at which KMTs are recruited, *j*(*s*). This relationship suggests a means to experimentally test models of KMT recruitment: since we directly measured *n*(*s*) by electron microscopy (i.e. Figure 6B), postulating a form *j*(*s*) allows *v*(*s*) to be calculated. The predicted *v*(*s*) can then be compared with measured KMT movements (Figure 3) to determine the extent to which it, and thus the postulated *j*(*s*), are consistent with both the electron microscopy and photoconversion data. This prediction requires specifying the rate of KMT detachment, which, based on our photoconversion measurements, we take to be *r* = 0.4 min^-1^.

We consider two models of KMT recruitment that have previously been proposed, either that KMTs are nucleated at kinetochores (Witt et al. 1980, Mitchinson and Kirschner 1985a, Khodjakov et al. 2000, Khodjakov et al. 2003, Maiatio et al. 2004, Sikirzhytski et al. 2018) or that KMTs arise from non-KMTs whose plus ends are captured by kinetochores (Mitchinson and Kirschner 1984, 1985b, 1986, Huitorel and Kirschner 1988, Rider and Alexander 1990, Hayden et al. 1990, Kamasaki et al. 2013, David et al. 2019). If all KMTs were nucleated at kinetochores, then *j*(*s*) = 0 everywhere in the spindle bulk (Figure 6D, upper). These “kinetochore-nucleated” KMTs could either be nucleated by the kinetochore itself or could be nucleated nearby and captured while still near zero length (Sikirzhytski et al. 2018). If instead all KMTs result from the capture of non-KMTs, then *j*(*s*) would be non-zero in the spindle bulk (Figure 6d, lower). In this latter case, *j*(*s*) would be the rate that a non-KMT whose minus end is at a position *s* along a streamline has its plus end captured by a kinetochore. We considered a model of non-KMT capture where any non-KMT can be captured provided that it reaches the kinetochore. We took the distribution of non-KMT minus ends along streamlines (Figure 6s2) as a proxy for the non-KMT nucleation rate, implying that *j*(*s*) is proportional to non-KMT minus end density times that probability that a nucleated non-KMTs grows long enough to reach the kinetochore before undergoing catastrophe and depolymerizing (see supplement). The kinetochore nucleation model predicts that the minus end speed monotonically increases with distance away from the pole along streamlines (Figure 6E). The non-KMT capture model predicts that the speed is near zero throughout the spindle. The two models thus offer qualitatively different predictions for KMT motions.

To understand why the two models offer qualitatively different predictions for the KMT minus end speeds, it is helpful to consider the contribution of each of the terms in the mass conservation, Equation 1, separately in the spindle bulk, where the minus end density distribution is roughly flat. In the nucleate at kinetochore model, the recruitment term, *j*(*s*), is zero by definition. The first KMT minus end motion flux term 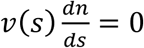 as well because the minus end density distribution is flat (i.e. 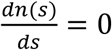). This leaves only the second KMT minus end motion flux term, 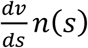, which describes changing KMT minus end speed and the detachment term *rn*(*s*), giving 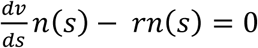, or equivalently 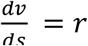. Thus, a linear increase in the speed of the KMTs with distance from the pole balances the constant detachment term in the spindle bulk. In contrast, in the capture from spindle model, the *j*(*s*) recruitment term is non-zero and can counteract the detachment terms in place of the changing speed term. The experimentally observed density of non-KMT minus ends is roughly the same as the density of KMT minus end along streamlines, so the newly nucleated KMTs roughly recapitulate the observed distribution, leaving a near-zero speed everywhere in the capture from spindle model. Therefore, the nucleate at kinetochore model predicts that the speed of KMT minus ends will increase with distance from the pole while the capture from spindle model predicts the KMT minus end speed is near-zero throughout the spindle.

### A simulation of the photoconversion experiment with nucleation at the kinetochore is consistent with the observed speed of tubulin

We next sought to determine whether the predictions from either the nucleate at kinetochore model or the capture from the spindle model were consistent with the motions of tubulin measured from photoconversion experiments. To do so, we simulated the motion of a photoconverted line of tubulin in the spindle using the two different models for KMT recruitment with the dynamics inferred from the flux balance analysis (Figure 6E).

Our simulations used a discrete model of KMTs with recruitment, growth, and detachment along streamlines in the spindle. At each timestep of the simulation, we generated newly recruit KMTs with Poisson statistics. The plus end position of these new KMTs was selected from the experimentally measured density distribution of kinetochores along streamlines (binned from all three reconstructed spindles) (Figure 7s1). The initial position of the minus ends of these new KMTs depended on the recruitment model: for the kinetochore nucleation model, the KMT minus end started at the position of kinetochores; in the capture from spindle model, the initial KMT minus end position was drawn from the (non-zero) distribution *j*(*s*) (see supplement). Thus, in the kinetochore nucleation model, newly recruited KMTs start with zero length (since they are nucleated at kinetochores), while in the spindle-capture model KMTs begin with finite length (since they arise from non-KMTs whose plus ends bind kinetochores). After a lifetime drawn from an exponential distribution with a detachment rate *r* = 0.4 min^-1^ (based on our photoconversion measurements), the KMT detaches from the kinetochore and is removed from the simulation.

In our model, newly polymerized tubulin incorporates at stationary, kinetochore bound KMT plus ends, while their minus ends move backwards along the streamline towards the pole with the experimentally inferred speed *v*(*s*), which varies based on the recruitment model (Figure 6E). In the absence of minus end depolymerization, all of the tubulin in a KMT moves at the same speed as its minus end *v_tub_*(*s*) = *v*(*s*), for a KMT whose minus end is at position *s*. If, however, the minus end of a KMT depolymerizes with a speed *v_tread_*(*s*), then the tubulin in the KMT will move faster than its minus end, at speed *v_tub_*(*s*) = *v*(*s*) + *v_tread_*(*s*). Based on a “chipper-feeder” model of minus end depolymerization, we included minus end depolymerases only at the spindle pole (Gabbe and Heald 2004, Dumont and Mitchinson 2004, Long et al. 2020). KMT minus ends in the spindle bulk thus move along streamlines without minus end depolymerization. When KMT minus ends enter the pole region at position *s_p_* = 1.5 *μm* along a streamline, the tubulin continues to incorporate at the plus end at the same speed as at the pole boundary, but minus end depolymerization begins, leading to tubulin to treadmill through the KMT at speed *v_tread_*(*s*) = [*v*(*s_p_*) − *v*(*s*)]θ(*s_p_* − *s*), where θ(*s*) is the Heavyside step function.

Both the kinetochore nucleation model and the capture from spindle model reproduce the experimentally measured KMT minus end distribution along streamlines (Figure 6S3), as they must by construction. We next considered a 2D slice of a spindle (to replicate confocal imaging) and modeled photoconverting a line of tubulin in the spindle with a modified Cauchy profile, which fits the shape of the experimentally converted region well (Figure 7S2). We simulated the motion of tubulin in individual KMTs and summed the contributions of each KMT together to produce a final simulated spindle image. Such simulations of the kinetochore nucleation model showed a steady poleward motion of the photoconverted tubulin (Figure 7A). In contrast, simulations of photoconverted tubulin in the capture from spindle model exhibited substantially less motion (Figure 7S3). To facilitate comparison to experiments, we analyzed the simulations with the same approach we used for photoconversion data. First, we projected the simulated photoconverted tubulin intensity onto the spindle axis to find the photoconverted line profile over time (Figure 7A, lower). We then fit the simulated line profile to a Gaussian and tracked the position of the peak over time to determine the speed of tubulin at the location of photoconversion. We varied the position of the simulated photoconversion line and repeated this procedure, to measure the speed of tubulin throughout the spindle in the two recruitment models (Figure 7B). The predicted spatially varying speeds of tubulin in the kinetochore nucleation model are consistent with experimentally measured values (Figure 3F), while the prediction from the capture from spindle model are too slow. If minus end depolymerization at the pole is turned off in the simulations, then the predicted speeds from both recruitment models become inconsistent with the experimental data (Figure 7S4).

**Figure 7:**
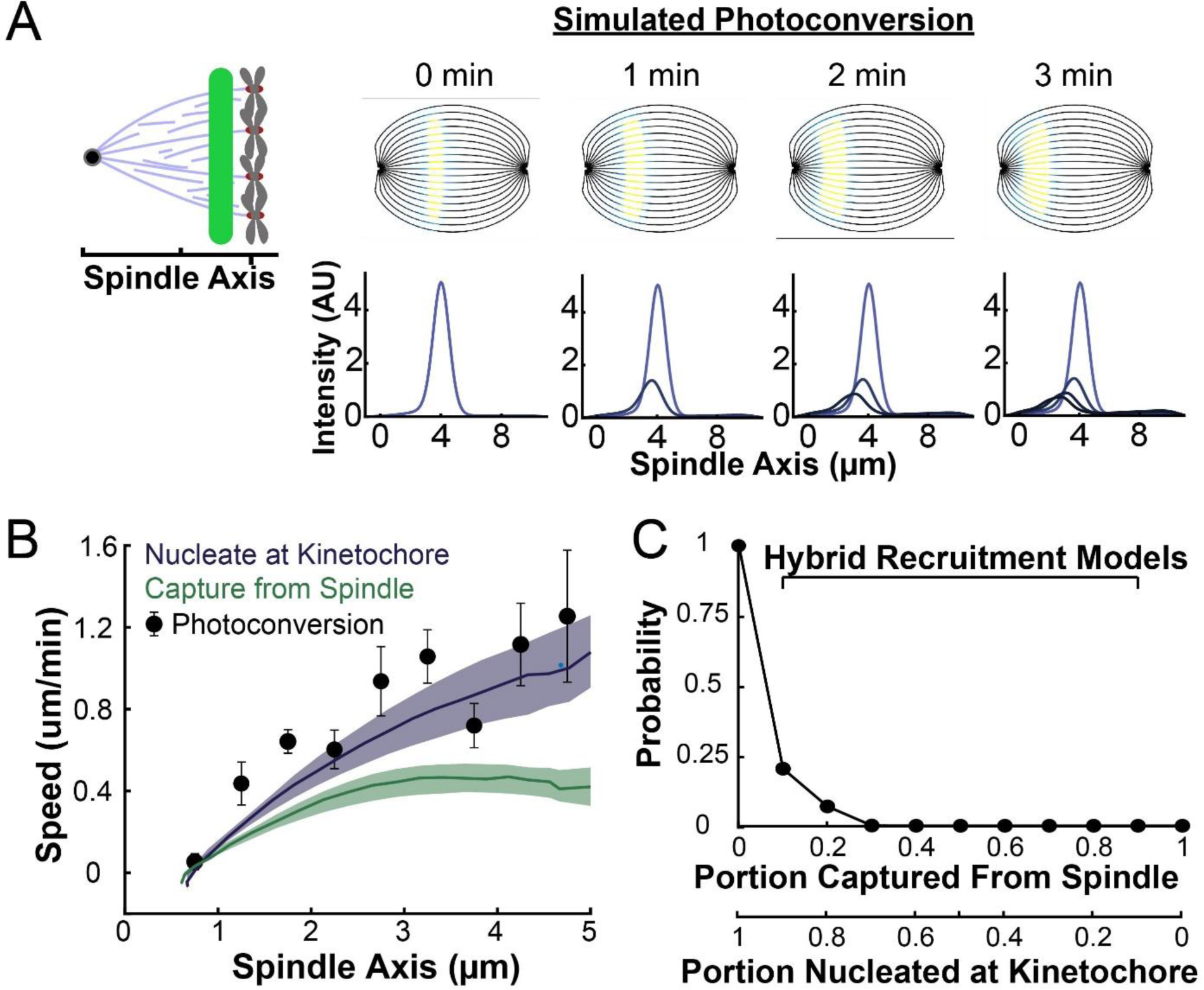
Model predicted tubulin flux compared to observed values. A) Sample simulated images and line profiles from a photoconversion simulation using KMT minus end speeds in the nucleate at kinetochore model. B) Comparison of the predicted spatial dependence tubulin flux speed in the nucleate at kinetochore and capture from spindle models. Error bars are standard error of the mean. C) Relative probabilities of hybrid version of the two models.

Our analysis showed that a model in which all KMTs nucleate at kinetochores is consistent with the observed speeds of tubulin throughout the spindle, while a model in which all KMTs are captured from the spindle bulk is inconsistent with this data. We next considered hybrid models which contained both KMT recruitment mechanisms. We simulated the motion of a line of photoconverted tubulin and varied the portion of KMTs nucleated at the kinetochore vs. captured from the spindle. We compare the feasibility of predictions from hybrid models with the data by calculating the Bayesian probability of observing the measured speeds with a uniform prior (Figure 7C). The model probability peaks at the edge where all KMTs are nucleated by the kinetochore. Thus, while the observed speeds are not inconsistent with a small fraction (less than 20%) of KMTs being captured from the spindle bulk, the data favors a model where KMTs are exclusively nucleated at kinetochores.

### A quantitative 3D model of KMT nucleation, minus end motion and detachment

We therefore propose a model where KMTs nucleate at kinetochores and grow along streamlines (Figure 8A). As the KMTs grow, they slow down until they reach the pole where minus end depolymerization causes tubulin to treadmill through the MT. The KMTs detach from the kinetochore at a constant rate, independent of their position in the spindle.

**Figure 8:**
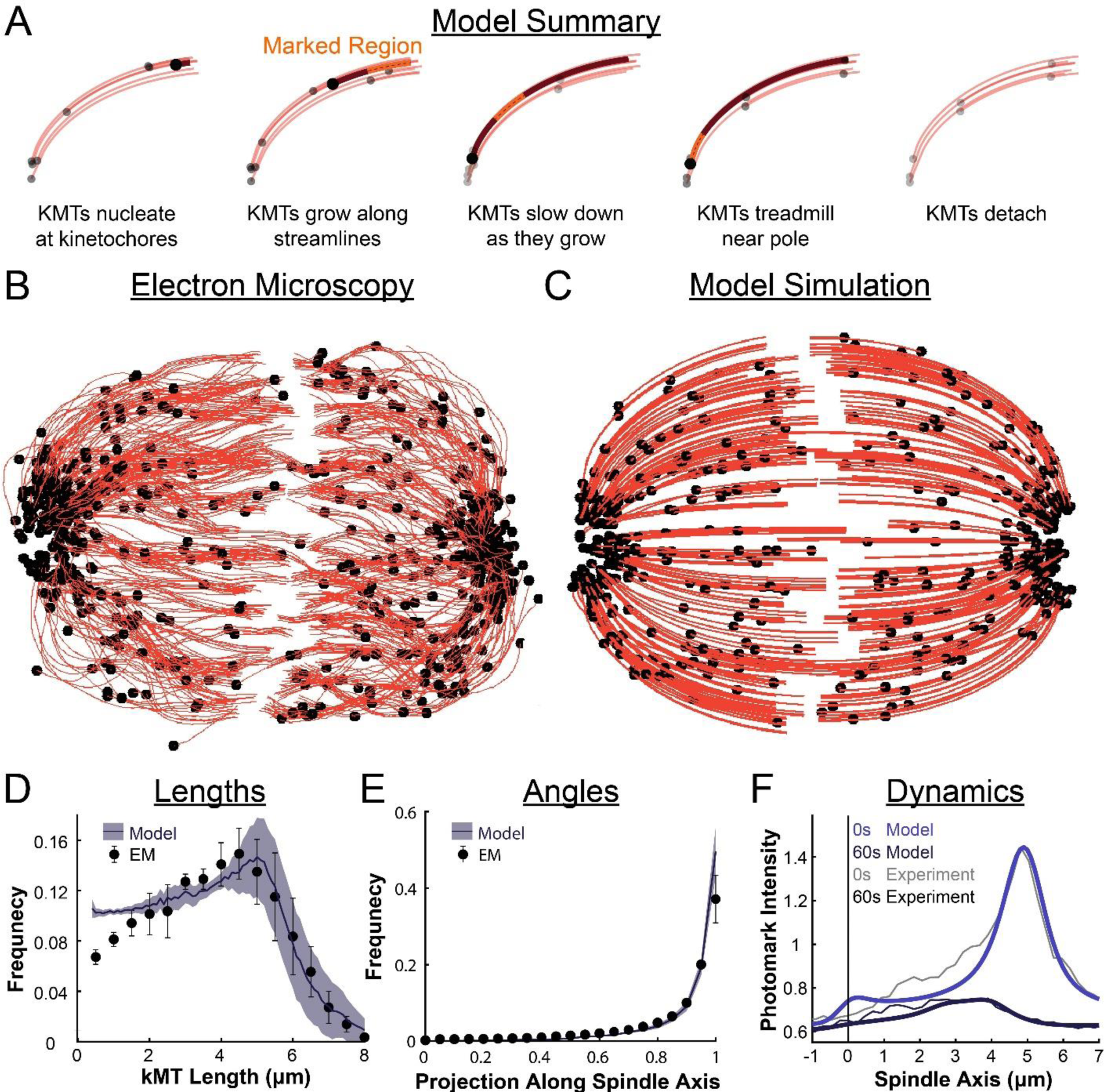
Summary of a nucleate at kinetochore model of KMT dynamics and structure in HeLa cells. A) Summary of the steps of the model: 1. KMTs nucleate at kinetochores 2. KMTs grow along stremalines 3. KMTs slow down as they grow 4. KMTs treadmill near the pole 5. KMTs detach. B) KMT structure from a sample EM reconstruction (Kiewisz et al., 2021; spindle #2). C) Model simulation of the KMT structure given the spindle geometry and kinetochore positions. D) Comparison of predicted and observed KMT lengths averaged over all three EM cells (Purple-model prediction, black EM data. Error bars are standard error of the mean). E) Comparison of predicted and observed KMT angles averaged (Purple-model prediction, black EM data. Error bars are standard error of the mean). F) Comparison of predicted and observed photoconverted line profiles (blue-model prediction, grey-experiment. Lighter shades are 0s, darker shades are 60s).

To test our model predictions against the full 3D reconstructed KMT ultrastructure of each spindle, we simulated the nucleation, growth and detachment of KMTs in 3D for each spindle separately. In each spindle, we simulated KMT nucleation by placing newly formed, zero length KMTs at the reconstructed kinetochore positions with Poisson statistics. The KMT minus ends then move towards the pole at the experimentally inferred speed *v*(*s*) undergo minus end depolymerization near the pole causing tubulin to treadmill at speed *v_tread_*(*s*) = [*v*(*s_p_*) − *v*(*s*)]θ(*s_p_* − *s*), and detach with a constant rate r.

The agreement between the electron tomography reconstruction (Figure 8B) and the predicted model structure is striking (Figure 8C, Video 8S1). We next compared the lengths of KMTs from the simulations with the experimentally measured length distribution. We found the lengths of KMTs in the simulated spindles by measuring the distance between the minus and plus end along the model KMT streamline trajectory; in the reconstructed spindles we traced the arclength of each KMT along its reconstructed trajectory. We binned the KMT lengths for each simulated and reconstructed spindle and averaged the spindles together to obtain the KMT length distributions. (Figure 8D). The observed length distribution of the KMTs from the reconstructed spindles is well predicted by the model. To compare the orientation of the simulated and reconstructed KMTs, we divided the MTs into short 100 nm sections and projected these sub-segments onto the spindle axis to find what portion of the section lie on the spindle axis. We binned the projections from each spindle and averaged the three resulting distributions together to obtain the distribution of projected lengths along the spindle axis (Figure 8E). There is similarly good agreement between the simulation prediction and the reconstructed projected lengths along the spindle axis. Both the predicted lengths and orientations of the KMTs are thus consistent with the ultrastructure measured by electron tomography.

We finally tested whether the predicted tubulin motion was consistent with the photoconversion experiment. We simulated the motion of a photoconverted plane of tubulin (with a modified Cauchy intensity profile) as we did in the 2D confocal case, but now moved the tubulin along 3D nematic trajectories (Movie 8S1). To simulate confocal imaging, we projected a thin 1μm confocal z-slice centered at the poles onto the spindle axis over the course of the simulation to produce a line profile. The simulated line profile agrees well with the experimental profile even after 60 seconds of simulation time (Figure 8F), indicating that the dynamics of the model are consistent with the experimentally measured tubulin motion and turnover. Taken together, these results favor a model where KMTs nucleate at the kinetochore, grow and slow down along nematic streamlines, undergo minus end depolymerzation near the pole and detach with a constant rate. Such a model is consistent with both measuremnts of KMT ultrastructure, from EM, and measurements KMT dynamics, from photoconversion, in HeLa cells.

## DISCUSSION

In this study, we leveraged recent electron tomography reconstructions that contain the positions, lengths, and configurations of microtubules in metaphase spindles in HeLa cells (Kiewisz et al. 2021). We used this dataset, in combination with live cell microscopy measurements and biophysical modeling, to investigate the behaviors of KMTs. We found that roughly half of KMT minus ends were not located at the poles (Figure 1). To better understand this KMT minus end distribution we performed a series of photoconversion experiments to measure the dynamics of KMTs. The fraction of slow turnover tubulin measured from photoconversion matched the fraction of tubulin in KMTs measured by electron tomography. This observation argues that KMTs are the only MTs in metaphase spindles that are appreciably stabilized. The photoconversion experiments also showed that the speed of tubulin in KMTs slowed down near the poles and that KMT turnover was uniform throughout the spindle (Figure 2 and 3). We found that both KMTs and non-KMTs were highly aligned (Figure 4) and that the orientations of MTs throughout the spindle can be quantitively explained by an active liquid crustal theory in which MTs locally align with each other due to their mutual interactions (Figure 5). This suggests that KMTs tend to move along well-defined trajectories in the spindle, so we analyzed the distribution of KMT minus ends along these trajectories (Figure 6). From these distributions, we predicted the speed of KMT minus ends using a mass conservation analysis. This analysis depends on the model of how KMTs are recruited to the kinetochore. We found that predictions from the nucleate at kinetochore model agreed well with the experimental measurements while the predictions from the capture from spindle model did not (Figure 7). We therefore propose a model where KMTs are nucleated at the kinetochore and polymerize from their plus ends as their minus ends move backwards along nematic streamline trajectories towards the pole, slowing down as they approach the pole. KMTs detach from the kinetochore at a constant rate. This model accurately predicts the lengths, orientations, and dynamics of KMTs in mitotic spindles of HeLa cells (Figure 8).

Previous work has shown that the photoconversion of tubulin in the spindle implies that there are at least two population of MTs, one with fast and one with slow turnover (Gorbsky and Borisy 1989, DeLuca et al. 2016, Warren et al. 2020). While the slow turnover fraction has often been ascribed to KMTs (Zhai et al 1995, Deluca et al 2010, Kabeche and Compton 2013), some work has suggested that substantial fractions of non-KMTs may be stabilized as well (Tripton 2021). We found that the fraction of tubulin in KMTs identified structural from the EM reconstructions (25±2%) and the stable fraction from the photoconversion experiments (24±2%) are statistically indistinguishable (Figure 2). This agreement argues that KMTs account for the overwhelming majority of stable MTs in the spindle. Thus, the slow decay rate can be interpreted as the rate of KMT turnover. Our observation that the slow decay rate is uniform thorough out the spindle, suggests that KMTs detached from the kinetochore at a constant rate, independent of the position of their minus ends in the spindle (Figure 3). The speed of a photoconverted line of tubulin is slower for lines drawn near the poles than in the center of the spindle. Since the speed of tubulin moving in KMTs in the spindle bulk is coupled to tubulin polymerization at the KMT plus end, this suggests that longer KMTs grow more slowly than shorter KMTs.

We found that MTs in spindles in HeLa cells were well-aligned with a high scalar nematic order parameter along orientations that are consistent with the predictions of an active liquid crystal theory. This implies that the orientations of MTs in the spindle is dictated by their tendency to locally align with each other. The tendency of MTs in the spindle to locally align with each other could result from the activity of MT crosslinkers, such as dynein, kinesin-5, or PRC1 (Kapitein et al. 2005, Tanenbaum et al. 2013, Wijeratne and Subramanian 2018), or simply from steric interactions between the densely packed rod-like MTs. The volume fraction of MTs in the reconstructed spindles is 0.052±0.05, which is slightly above the volume fraction where the nematic phase is expected to become more stable than the isotropic phase (∼0.04) (Doi and Edwards 1988. Brugués and Needleman 2014). Steric interaction between the MTs could therefore be enough to explain the observed nematic behavior. Studying spindles with depleted crosslinking proteins, lower MT density and perturbed KMT dynamics would help to determine the origin of these aligning interactions.

It has previously been unclear to what extent KMTs nucleate de novo at kinetochores vs resulting from non-KMTs being captured by the kinetochore (Tezlzer et al. 1975, Mitchinson and Kirschner 1985a, Mitchinson and Kirschner 1985b, Huitorel and Kirschner 1988, Heald and Khodjakov 2015, LaFountain and Oldenborug 2014, Petry 2016, Sikirzhyski et al. 2018, David et al. 2019, Renda and Khodjakov 2021). We show that a model where KMTs nucleate at kinetochores is consistent with the KMT ultrastructure observed in the tomography reconstructions and the tubulin dynamics observed in the photoconversion experiments. Our results would also be consistent with a model in which specifically MTs nucleate very near the kinetochore and are rapidly captured.. Such a capture of short MTs near the kinetochore could be consistent with observations of short MTs near chromosomes during prometaphase (Sikirzhytski et al. 2018).

The present work combined large-scale EM reconstructions, light microscopy, and theory to study the behaviors of KMTs in metaphase spindles. The behaviors of KMTs may be dominated by other processes at those different times. In the future, it would be interesting to apply a similar methodology to investigate the behavior of KMTs during spindle assembly in prometaphase and chromosome segregation in anaphase. Another interesting direction would be to apply a similar methodology to the study of spindles in other organisms. Previous EM reconstructions in *C. elegans* mitotic spindles have found a similar distribution of KMT lengths in metaphase (Redemann et al 2017). Acquiring electron tomography reconstructions and dynamics measurements in a different model systems would help elucidate whether the proposed KMT lifecycle is conserved across metazoans or unique to human cells.

One significant feature of the nematic-aligned, nucleate-at-kinetochore model is that it provides a simple hypothesis for the mechanism of chromosomes biorientation: A pair of sister kinetochores, with each extending KMTs, will naturally biorient as the KMTs locally align along nematic streamlines that are flat near the center of the spindle. Once bioriented, newly nucleated KMTs from either sister will naturally grow towards opposite poles. Microtubules attached to the incorrect pole will turnover over and be replaced by newly nucleated microtubules that will integrate into the nematic network, growing towards the correct pole. Once all of the incorrect microtubules have been cleared, tension generated across the opposite sisters will stabilize the existing, correct attachments. The nematic aligned, kinetochore-nucleated picture thus provides a self-organized physical explanation for chromosome bi-orientation and the correction of mitotic errors. It will be an exciting challenge for future work to test the validity of this picture.

## MATERIALS AND METHODS

### HeLa Cell Culture and Cell Line Generation

HeLa Kyoto cells were thawed from aliquots and cultured in DMEM (ThermoFisher) supplemented with 10% FBS (ThermoFisher) and Pen-Strep (ThermoFisher) at 37°C in a humidified incubator with 5% CO_2_. Cells were regularly tested for mycoplasma contamination (Southern Biotech).

Three stable HeLa cell lines were generated using a retroviral system. A stable HeLa Kyoto cell line expressing mEOS3.2-alpha tubulin and CENPA - GFP was generated and selected using puromycin and blasticidin (ThermoFisher) (Yu et al. 2019). An additional mEOS3.2-alpha tubulin and SNAP-Centrin cell line was generated and selected using puromycin, blasticidin and hygromycin. A final cell line expressing CENPA-GFP and GFP-Centrin was generated and selected using puromycin and hygromycin.

### Spinning Disc Confocal Microscopy and Photoconversion

All photoconversion experiments were performed on a home built spinning disc confocal microscope (Nikon Ti2000, Yokugawa CSU-X1) with 488nm, 561nm and 647nm lasers, an EMCCD camera (Hamamatsu) and a 60x oil immersion objective. Imaging was controlled using a custom Labview program (Wu et al. 2016). Two fluorescence channels were acquired every 5s with either 300ms 488nm and 500ms 561nm exposure for the initial imaging with the pre-converted frame or with 500ms 561nm and 300ms 647nm exposure for experiments with the SNAP-Centrin pole marker. The mEOS3.2 was photoconverted using a 405nm diode laser (Thorlabs) and a PI-XYZnano piezo (P-545 PInano XYZ; Physik Instrumente) to draw the photoconverted line. The line was moved at a speed of 5um/s with a laser power of 500nW (measured at the objective). Cells were plated onto 25-mm-diameter, #1.5-thickness, round coverglass coated with poly-d-lysine (GG-25-1.5-pdl, neuVitro) the day before experiments. Cells were stained with 500nM SNAP-SIR (New England Biolabs) in standard DMEM media for 30 minutes and then recovered in standard DMEM media for at least 4 hours. Before imaging, cells were pre-incubated in an imaging media containing Fluorobrite DMEM (ThermoFisher) supplemented with 10mM HEPES for ∼15min before being transferred to a custom-built cell-heater calibrated to 37°C. In the heater, cells were covered with 750μL of imaging media and 2.5mL of mineral oil. Samples were used for roughly 1 hour before being discarded.

### Quantitative Analysis of Photoconversion Data

All quantitative analysis was performed using a custom MATLAB GUI. We first fit the tracked both poles using the Kilfiol tracking algorithm (Gao and Kilfoil 2009) and defined the spindle axis as the line passing between the two pole markers. We generated a line profile along the spindle axis was then generated by averaging the intensity in 15 pixels on either side of the spindle axis. The activated peak from each frame was fit to a Gaussian using only the central [check number] pixels. If multiple peaks were identified, the peak closest to the peak from the previous frame was used. The position of the peak was defined to be the distance from the center of the peak to the pole marker. To determine the height of the peak, we subtracted the height of the gaussian from the height of a gaussian fit on the opposite side of the spindle to correct for background and divided by a bleaching calibration curve.

### Bleaching Calibration

HeLa spindles were activated by drawing 3 lines along the spindle axis from pole to pole. We then waited 5 minutes for the tubulin to equilibrate and began imaging using the same conditions as during the photoconversion measurement (561nm, 500ms exposure, 5s frames; 647nm, 300ms exposure, 5s frames). We calculated the mean intensity inside an ROI around the spindle (Figure 2s1a) and plotted the average of the relative intensity of 10 cells). We subtracted off a region outside of the cell to account for the dark noise of the camera. We then divide our intensity vs. time curve by the bleaching calibration curve to produce a bleaching-corrected intensity curve to fit to a dual-exponential model.

### Polarized Light Microscopy (PolScope)

We measured the orientation of spindle MTs in living cells using an LC-PolScope quantitative polarization microscope (Oldenbourg et al. 1998, Oldenbourg et al. 2005) The PolScope hardware (Cambridge Research Instruments) was mounted on a Nikon TE2000-E microscope equipped with a 100x NA 1.45 oil immersion objective lens. We controlled the PolScope hardware and analyze the images we obtained using the OpenPolScope software package. To ensure that the long axis of the spindle lies in or near the image plane, we labeled the poles with SNAP-Sir and imaged the poles using epifluorescence while we acquired the PolScope data. In all subsequent analysis, we use only data from cells where the poles lie within ∼ 1 μm of each other in the direction perpendicular to the image plane. To average the orientation fields from different spindles, we first determined the unique geometric transformation (rotation, translation, and rescaling) that aligns the poles. We then applied the same transformation to the orientation fields and took the average.

### Fitting Average MT Angles to Nematic Theory

For each 3D reconstructed spindle, the positions of the MTs were first projected into a 2D spindle axis-radial axis plane (averaging along the φ direction in cylindrical coordinates into a single plane, see Figure 4s3). Local MT angles were then averaged (<ϴ>=arg(<exp(2πiϴ)>)/2) in 0.1μm by 0.1μm bins in the spindle-radial plane.

We registered the three EM spindles by rescaling them along the spindle and radial axis. We rescaled the spindle axis of each spindle so all three spindles had the same pole-pole distance. We rescaled the radial axis so that the width of the spindles, measured by the width of an ellipse fit to the spindle density in the spindle axis-radial axis plane, was the same. We then averaged the three EM spindles together to produce Figure 5A. We similarly registered the PolScope images by rescaling the spindle axis using the pole-pole distance and the radial axis using the width of an ellipse fit to the spindle retardance image before averaging the cells together to produce Figure 5B.

The angles predicted by the active liquid-crystal model were found by solving the Laplace equation in the spindle bulk using a 2D finite difference method subjected to the tangential anchoring and defect boundary conditions. The model’s geometric parameters were determined by fitting the predicted angles to the averaged EM data by minimizing a χ^2^ statistic. We first fit the height, width and center of the elliptical boundary with the m=1 defects fixed at the edge using the averaged EM spindles. The elliptical boundary parameters were then fixed, and the position of the m=1 defects along the spindle axis and the radius of the defects were fit to produce Figure 5C. The individual spindles were similarly fit by first fitting the elliptical boundary with the m=1 defects on the edge and then fitting the position and radius of the defects to produce Figure 6s1.

| **Reagent type (species) or resource** | **Designation** | **Source of reference** | **Identifiers** | **Additional** |
| --- | --- | --- | --- | --- |
| Cell line (*Homo sapiens*) | HeLa Kyoto | Gerlich Lab, IMBA, Vienna Austria | - | - |
| Transfected construct (*Homo sapeins*) | pBABE-puro CENP-A:GFP | Yu et al. 2019 | - | CENP-A C-terminally labeled with sfGFP; in retroviral vector with puromycin selection marker |
| Transfected construct (*Homo sapeins*) | pBABE-hygro SNAP-Centrin | This paper | - | CENP-A C-terminally labeled with sfGFP; in retroviral vector with hygromycin selection marker |
| Transfected construct (*Homo sapeins*) | pJAG98(pBABE-blast) mEOS3.2-alpha tubulin | Yu et al. 2019 | - | CENP-A C-terminally labeled with sfGFP; in retroviral vector with blastcidin selection marker |
| Commerical assay or kit | SNAP-Cell 647-SiR | New England Biolabs | - | Catalog number S9102S |
| Software algorithm | Interactive spindle photoconversion analysis GUI (MATLAB 2020b) | This paper | - | - |
| Software algorithm | Photoconversion simulation package | This paper | - | - |
| Software algorithm | Photoconversion control and imaging | Wu et al. 2016 | - | Controls custom confocal photoconversion for arbitrary geometry |
| Software<br>algorithm | Polarizaed light<br>microscopy<br>control sofware | OpenPolScope.org | - | - |

## Supporting information

Video 8s1 - Simulated tubulin photoconversion in a 3D model spindle

## ACKNOWLEDGMENTS

The authors would like to thank Gloria Ha and Che-Hang Yu for technical assistance with experiments; Reza Farhadifar and Sebastian Fürthauer for helpful comments on the manuscript,. Research in the Needleman Lab is supported by the NSF-Simons Center for Mathematical and Statistical Analysis of Biology at Harvard (award number #1764269), and the Harvard Quantitative Biology Initiative. Will Conway was support by an NSF GRFP fellowship and an NSF-Simons Harvard Quantitative Biology Initiative student fellowship. Research in the Müller-Reichert laboratory is supported by funds from the Deutsche Forschungsgemeinschaft (MU 1423/8-2). R.K. received funding from the European Union’s Horizon 2020 research and innovation program under the Marie Skłodowska-Curie grant agreement No. 675737 (grant to T.M.R.).

---

**Figure 2s1:**
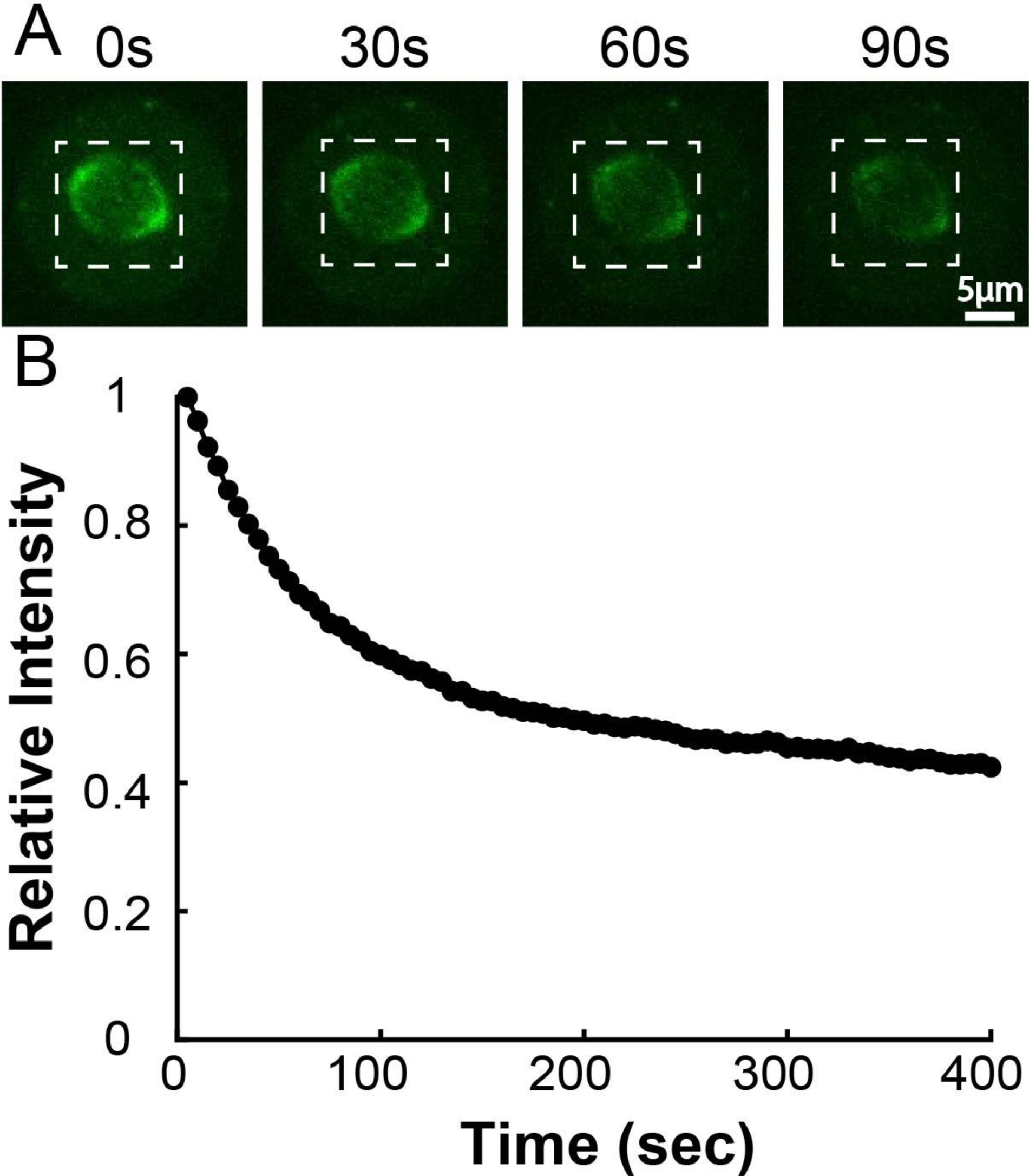
mEOS3.2-Alpha Tubulin Bleaching Calibration. A) Time series of activated tubulin in spindles. The whole spindle was photoactivated with 405nm UV light and left to equilibrate for 5 minutes before imaging B) Mean integrated spindle intensity over time (boxed region). Curves were corrected for dark background by subtracting the mean intensity of a small region marked outside the cell. Curves from 5 cells were normalized to the initial intensity at t=0s and then averaged together.

**Figure 4s1:**
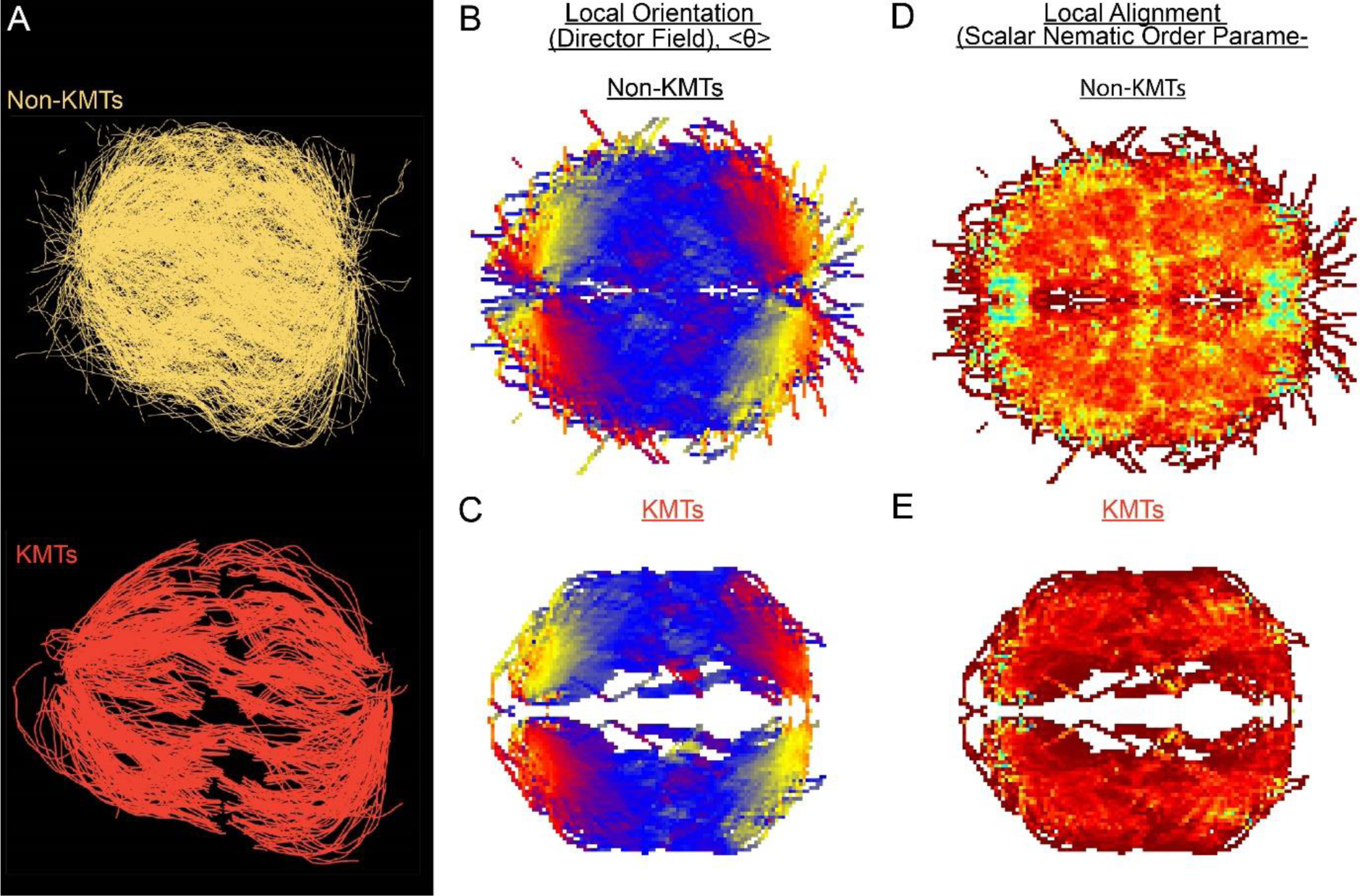
Measuring nematic alignment of non-KMTs and KMTs (reconstructed cell #2). A) Sample reconstruction from a single EM reconstruction of non-KMTs (yellow) and KMTS (red). B) Mean local orientation of non-KMTs average over all theta along the spindle axis. C) Mean local orientation of KMTs average over all theta along the spindle axis. D) Local alignment of the non-KMTs. E) Local alignment of the KMTs.

**Figure 4s2:**
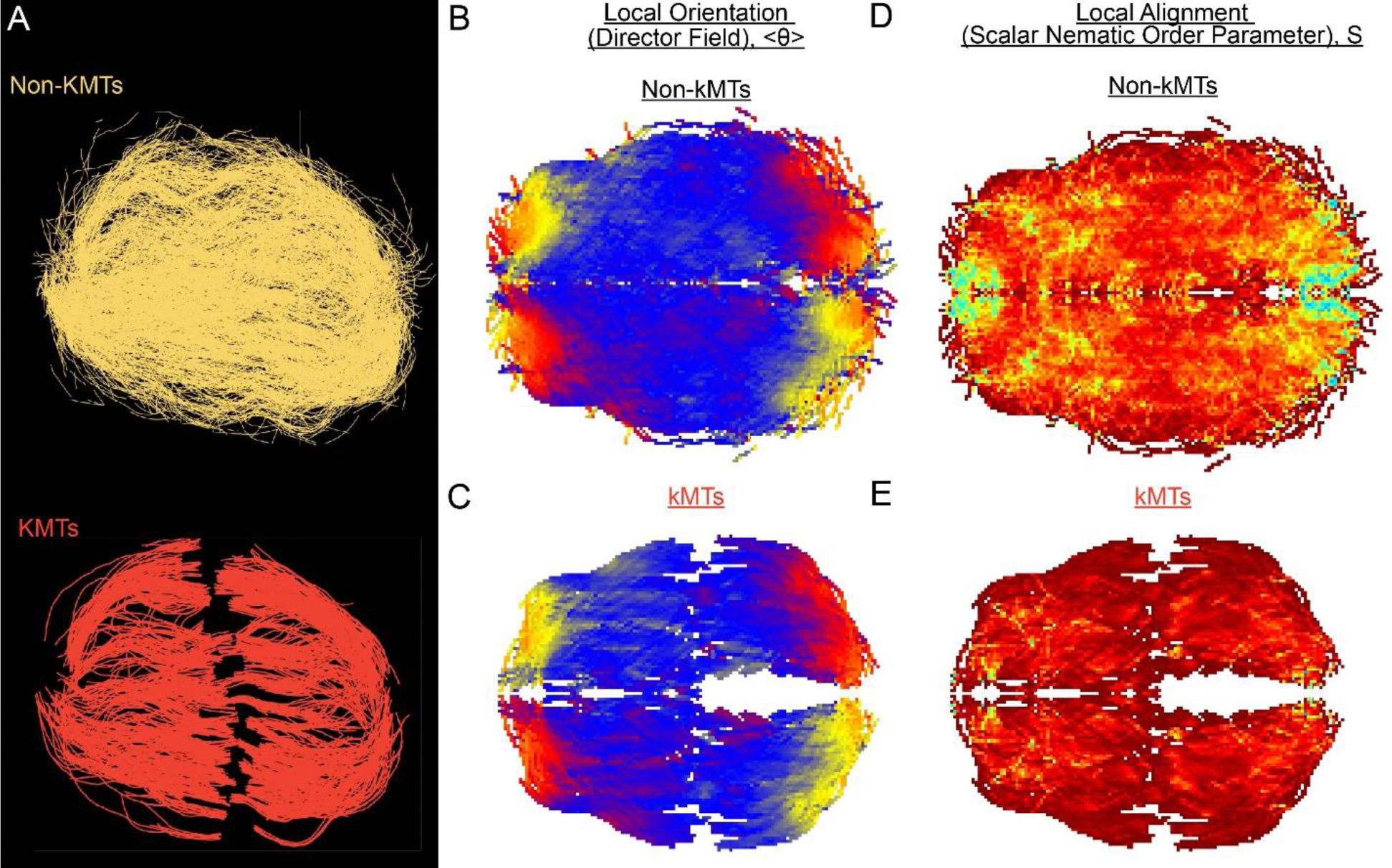
Measuring nematic alignment of non-KMTs and KMTs (reconstructed cell #3). A) Sample reconstruction from a single EM reconstruction of non-KMTs (yellow) and KMTS (red). B) Mean local orientation of non-KMTs average over all theta along the spindle axis. C) Mean local orientation of KMTs average over all theta along the spindle axis. D) Local alignment of the non-KMTs. E) Local alignment of the KMTs.

**Figure 4s3:**
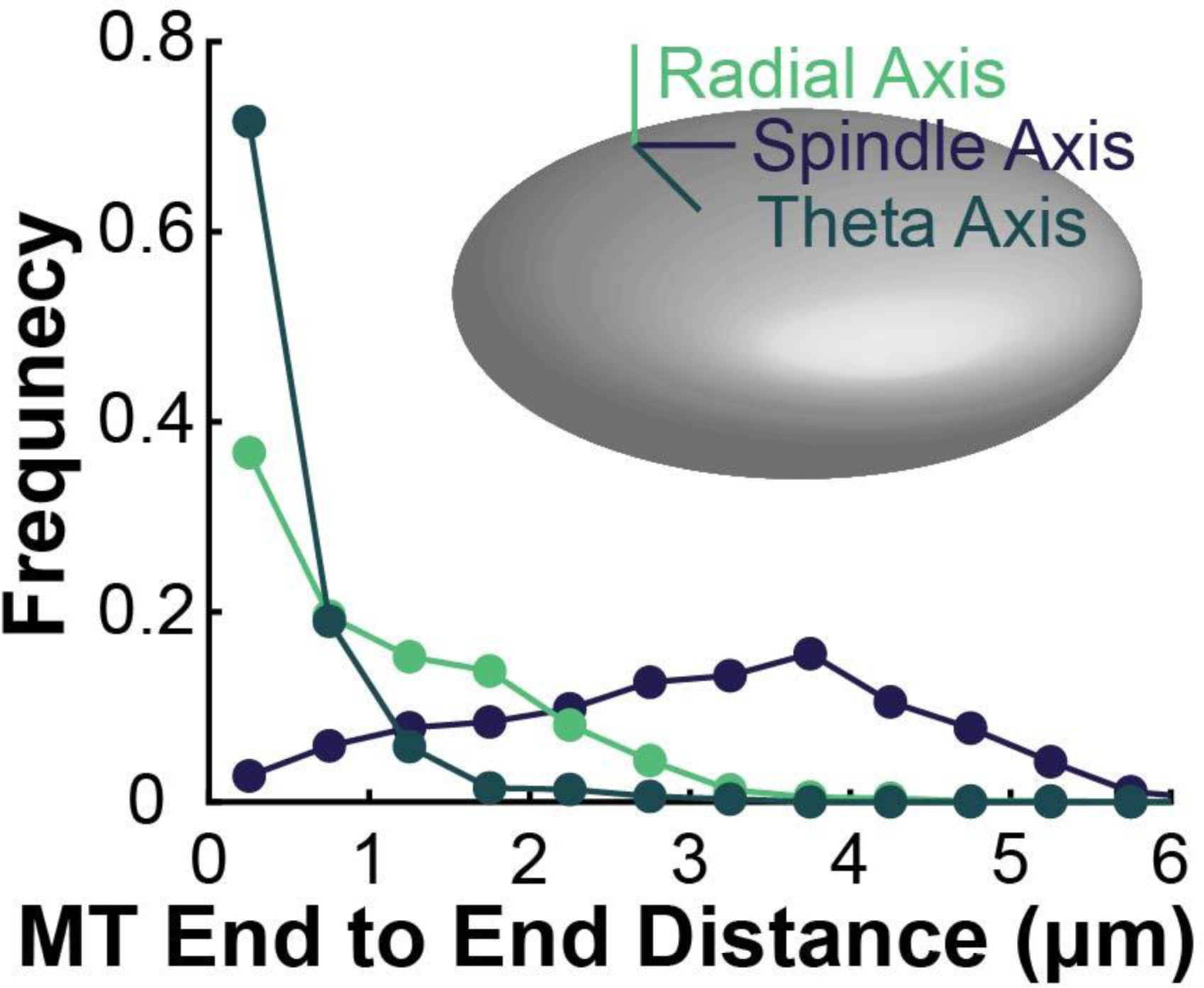
Microtubule end to end distance along the radial, spindle and theta axis. For each KMT, the distance between the plus and minus end of the microtubule along the radial (green), spindle (purple) and theta (cyan) axes and binned as a histogram. The radial, spindle, and theta axis are defined on the cartoon inset.

**Figure 5s1:**
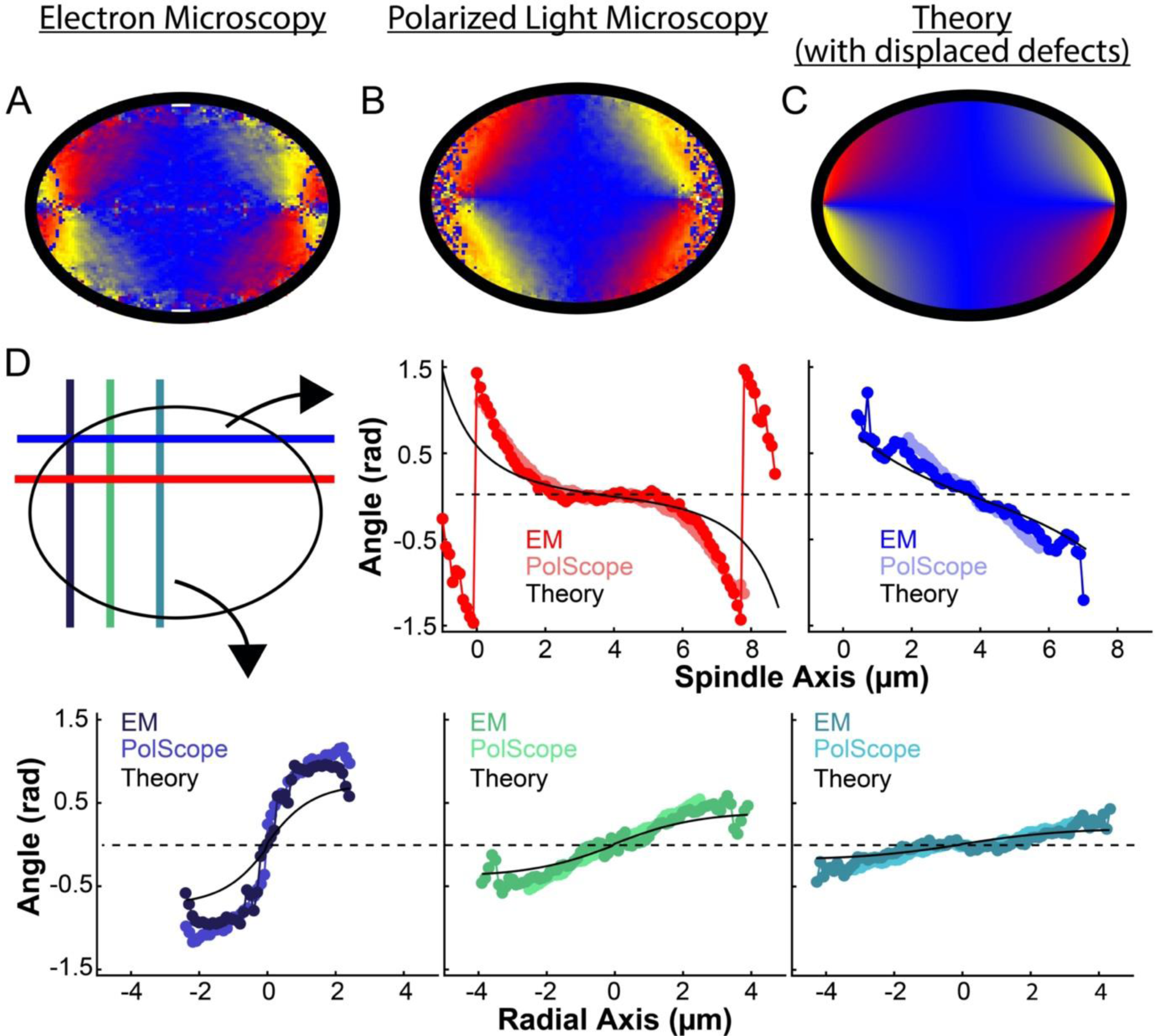
Experimentally measured orientation field of MTs in HeLa spindles compared to theoretical predictions with point defects localized on the spindle periphery. A) Orientation field of MTs from averaging EM reconstructions from three spindles. B) Orientation field of MTs from averaging polarized light microscopy (LC-PolScope) data from eleven spindles. C) A theoretical model of the spindle geometry with tangential anchoring at the elliptical spindle boundary and point defects on the spindle periphery. D) Average angle along narrow cuts parallel to the spindle and radial axis (red-lower spindle cut, blue-upper spindle cut purple-radial cut near pole, green-radial cut halfway between pole and kinetochore, teal-radial cut near kinetochore).

**Figure 6s1:**
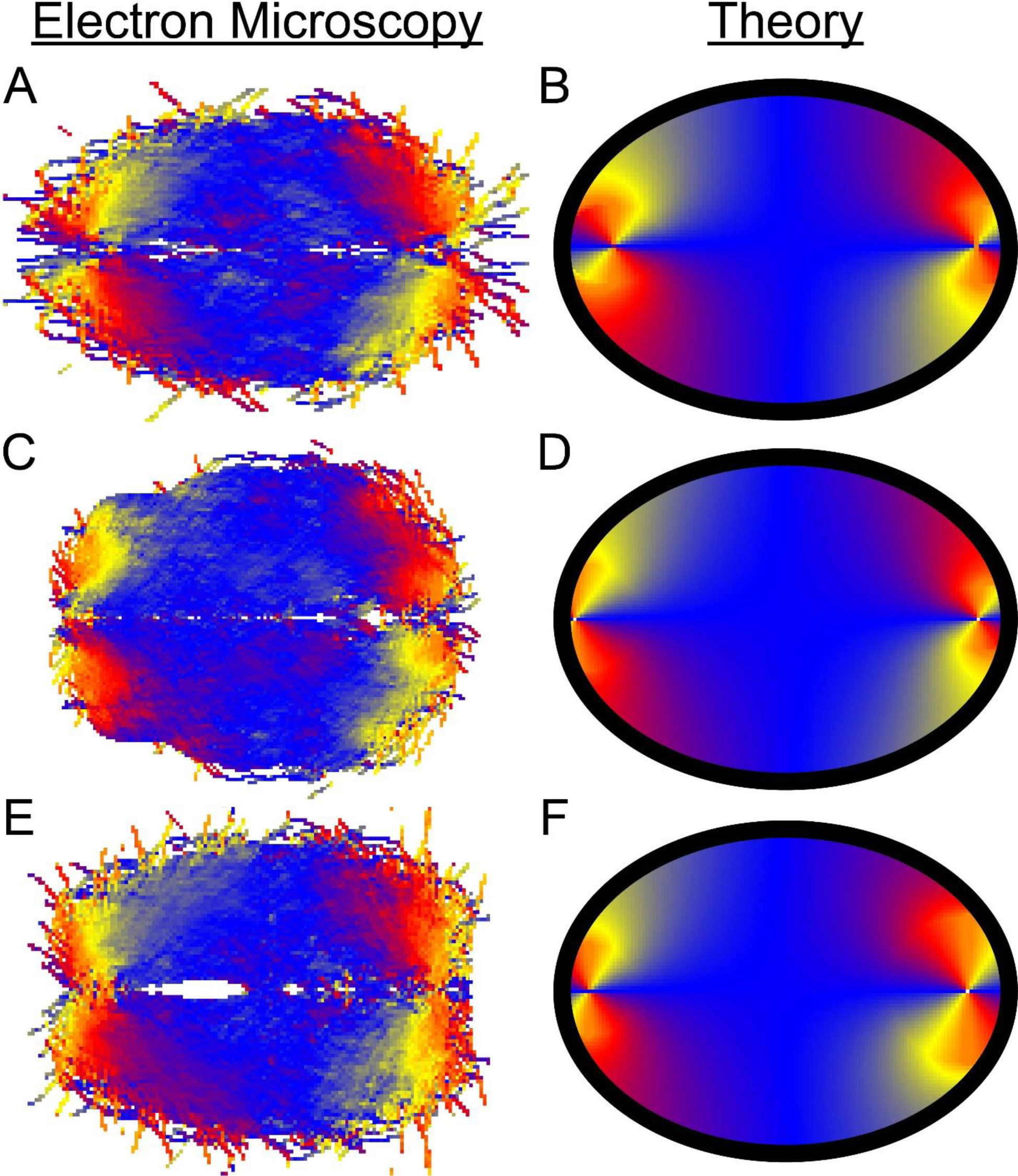
Comparison of EM and fit liquid crystal theory for individual reconstructed spindles. A) Average MT orientation from reconstructed spindle #1. B) Theoretical model of the spindle geometry with tangential anchoring at the elliptical spindle boundary conditions and point defects at the poles for spindle #1. C) EM for spindle #2. D) Theory for Spindle #2. E) EM for spindle #3. F) Theory for spindle #3.

**Figure 6s2:**
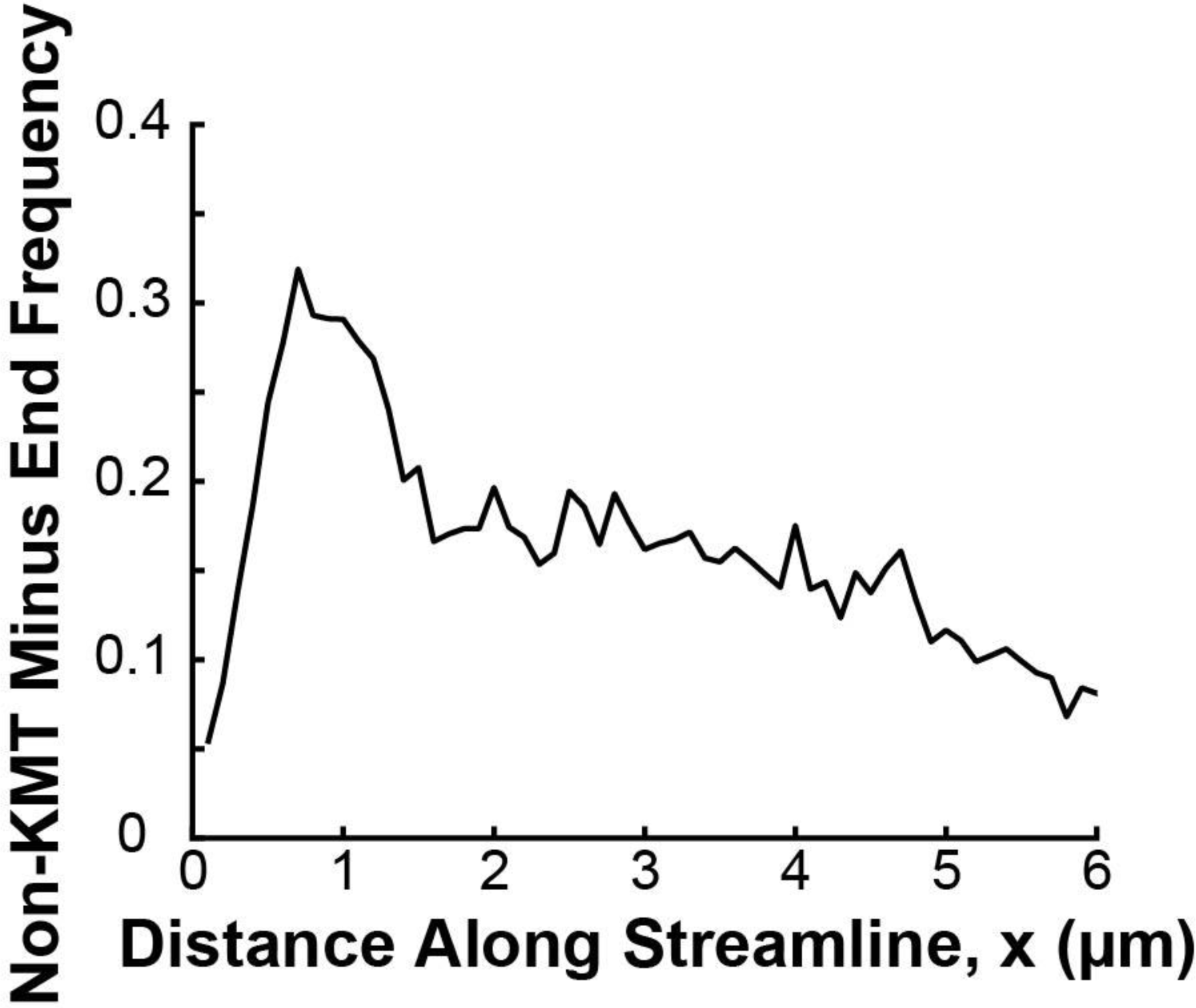
Density distribution of non-KMT minus ends along streamlines. For both ends of each non-KMT, the streamline trajectory from the non-KMT end was calculated by integrating along the nematic director field for that spindle. The distance from each end to the closer pole was then calculated, and the end closer to either pole was takem to be the minus end. The result from all three reconstructed spindles is shown in black.

**Figure 6s3:**
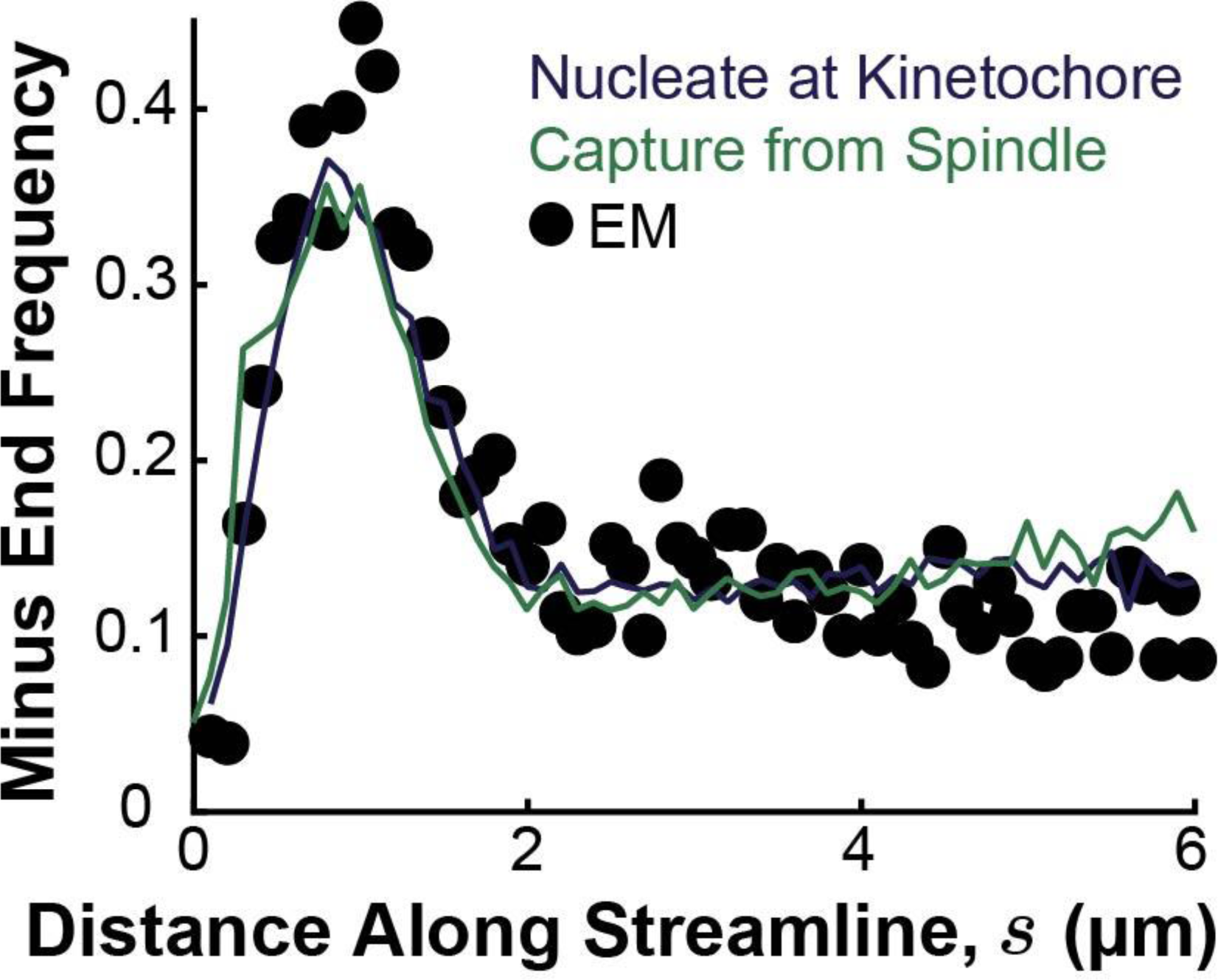
Simulated distribution of minus ends along streamlines using either a nucleate at kinetochore model (blue) or a capture from spindle recruitment model (green), compared to the experimentally measured minus distribution from electron microscopy reconstructions (black). KMTs were nucleated and plus ends were placed at positions drawn from the distribution of kinetochores along streamlines. For the capture from spindle model, the KMT minus ends were initially placed along streamlines at positions drawn from the distribution of tubulin density (from both KMTs and non-KMTs) along streamlines. For the nucleate at kinetochore model, KMT minus ends were placed at the kinetochore position. Minus ends were then moved along streamlines according to the velocities compute by either model until equilibrated to steady state.

**Figure 7s1:**
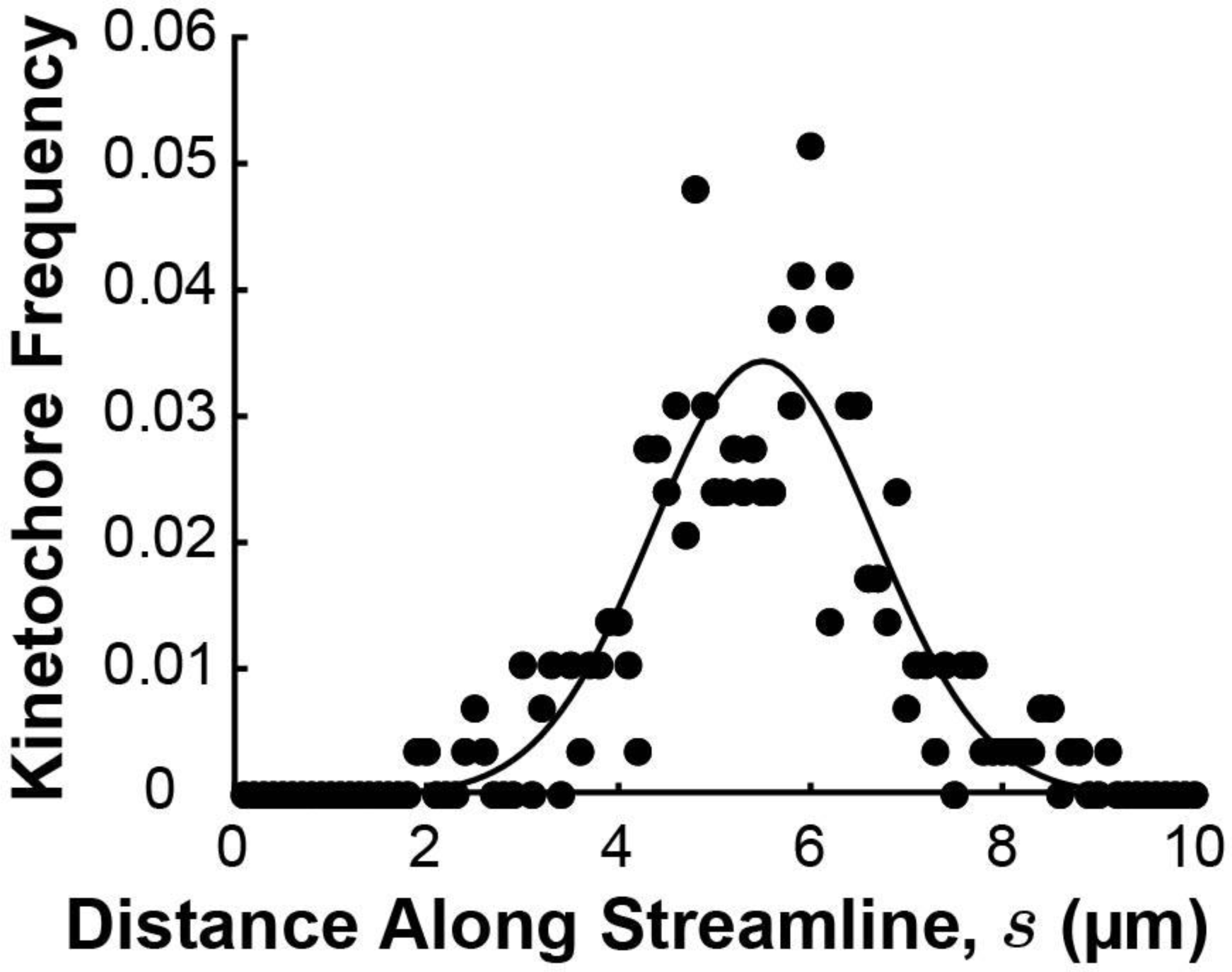
Density distribution of kinetochores along streamlines. The position of kinetochores in each sample cell was projected onto the streamline trajectories computed in figure 6s3 (black dots) and binned from all three cells. The experimental distribution was fit ot a Gaussian profile (solid black line)

**Figure 7s2:**
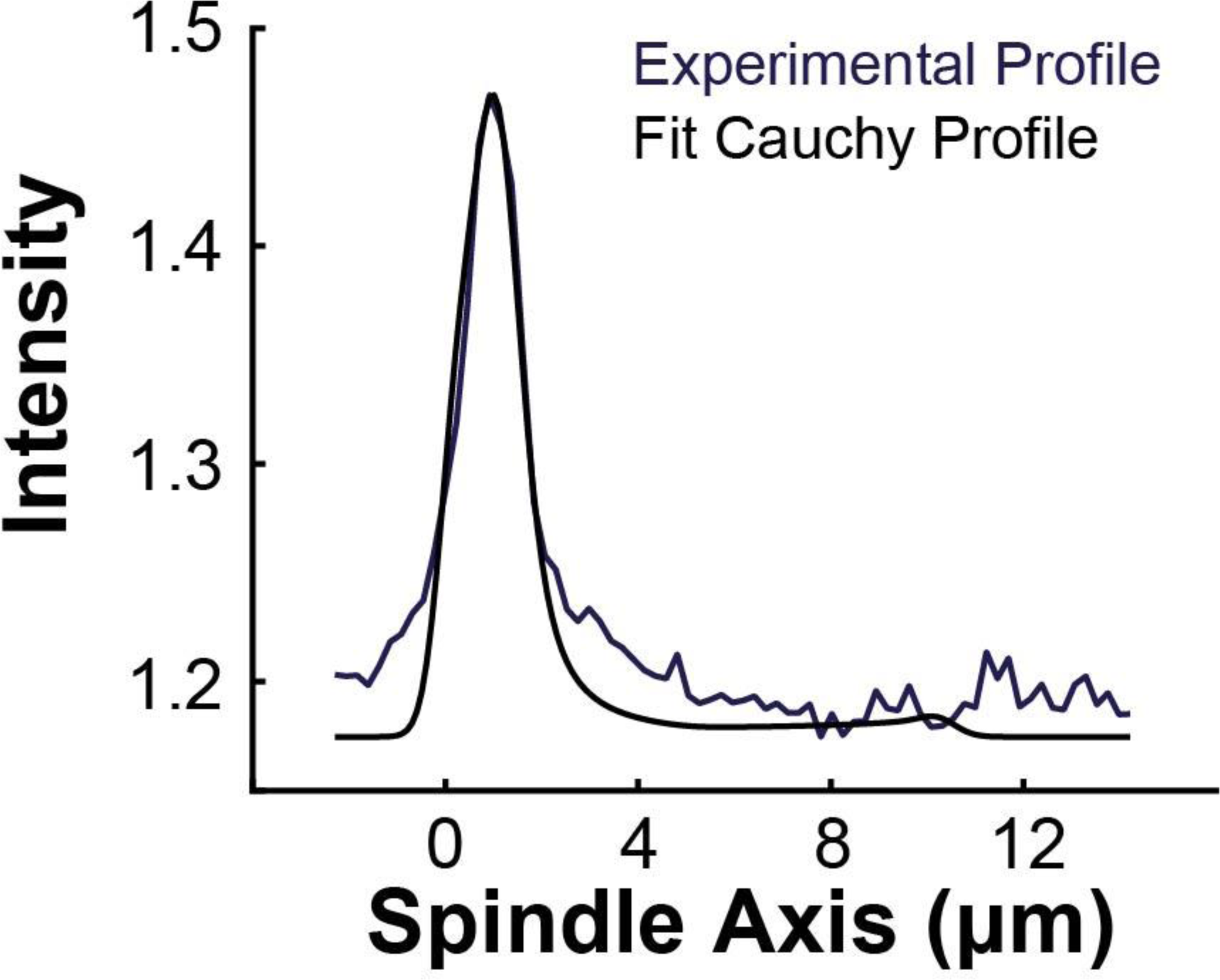
Sample experimental line profile from a photoconversion experiment and a fit modified Cauchy profile 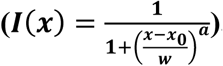, The fit profile was generated by drawing a photoconverted line on the simulated spindle (Figure 7A) and projecting the calculated tubulin intensity onto the spindle axis with the modified Cauchy profile with various central positions l0, widths w, and Cauchy exponent a. Best fit was determined from a χ^2^ minization algorithm.

**Figure 7s3:**
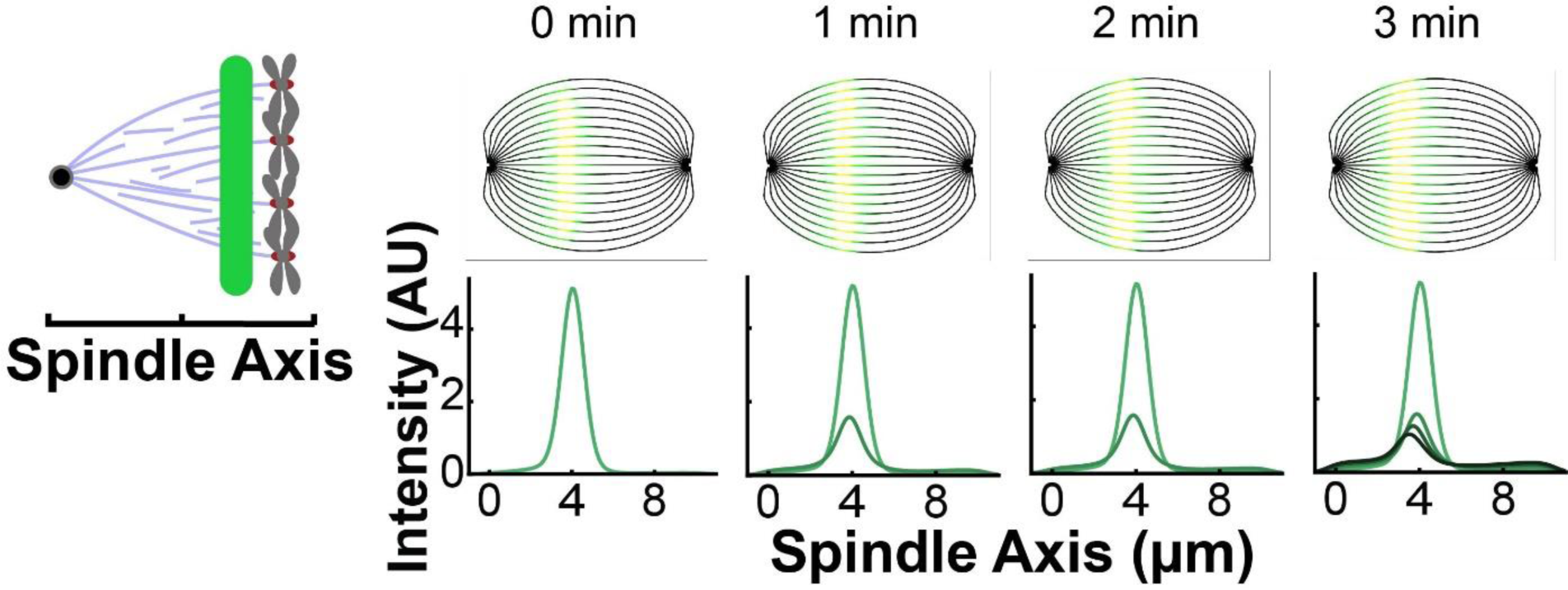
Sample simulated images from photoconversion in the capture from spindle model. For each streamline, a photoconverted line was drawn on the simulated, idealized spindle using the fit modified Caucy profile from Figure 7s2. The photoconverted tubulin intensity was then projected onto the spindle axis and summed across every streamline. The motion of the photoconverted tubulin along streamlines was calculated using the velocities from the capture from spindle model and is shown at subseuqenct times (30s, 60s and 90s).

**Figure 7s4:**
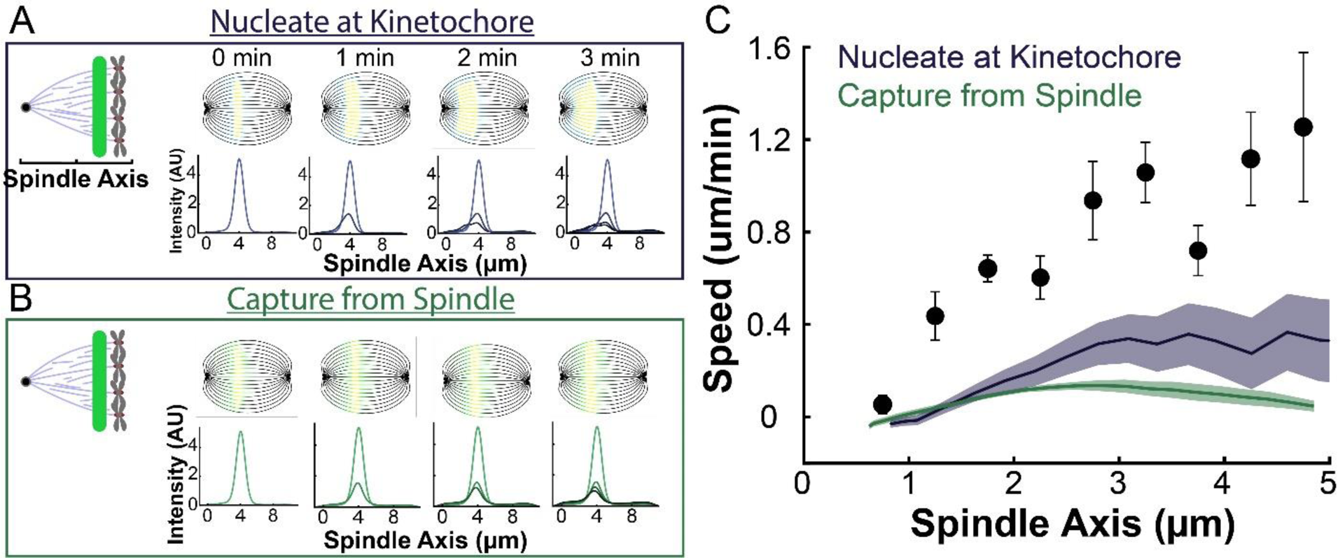
Model predicted tubulin flux compared to observed values without minus end depolymerization at the pole. A) Sample simulated images and line profiles from a photoconversion simulation using KMT minus end speeds in the nucleate at kinetochore model. B) Sample simulated images and line profiles from a photoconversion simulation using KMT minus end speeds in the capture from spindle model. C) Comparison of the predicted spatial dependence tubulin speed in the nucleate at kinetochore and capture from spindle models. Error bars are standard error of the mean.

**Movie 8s1:**
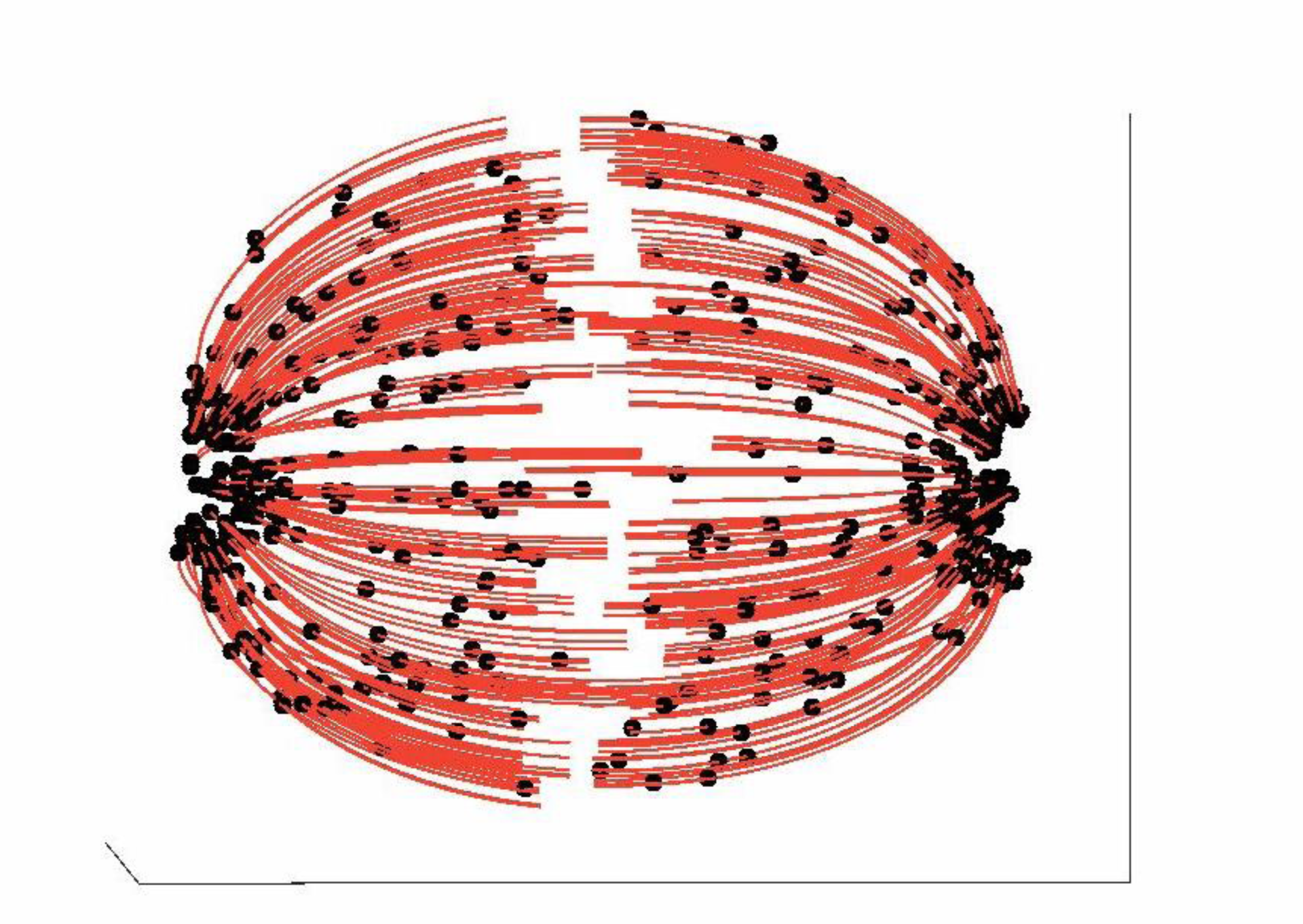
Simulated tubulin photoconversion in a 3D model spindle. Model simulation of the motion of motion of KMTs in a nucleate at kinetochore model. KMTs are shown in red, KMT minus ends are shown in black, photoconverted tubulin is shown in yellow. The model runs for 5 minutes of simulation time before the photoconverted line is drawn.

## Appendix 1: Computational Modeling Supplement

Here, we describe the details of the analysis, biophysical modeling, and simulations we performed to connect the structure of individual microtubules measured by electron tomography to the dynamics we observed in the photoconversion experiment. We first define the geometry of the simulated spindles. We then describe the details of the minus end speed prediction calculation and the simulation.

### Simulation spindle geometry

To generate idealized versions of each of the three reconstructed spindles for the simulations, we first separately fit each of the three spindles that were reconstructed by electron microscopy (EM) to an ellipse (Figure 6s1). We then fit the position and size of m=1 liquid crystal defects to the director fields of each spindle with tangential anchoring at the elliptical boundary (see Methods). In the simulations, we considered the motion of photoconverted tubulin along discrete nematic streamlines. We placed these streamlines 0.5µm apart at the center of the spindle along the radial axis (Figure A1). We found the trajectories of the streamlines by integrating along the director field predicted by a nematic model with tangential anchoring along the elliptical boundary and m=1 defects at the poles.

### Measuring the minus end density distribution *n*(*s*) from the EM reconstructed spindles

To measure the minus end density distribution *n*(*s*) along streamlines, we first found the position of every kinetochore microtubule (KMT) minus end along the fit nematic streamlines in each of the three EM reconstructions. For each KMT minus end, we found the streamline it was on by integrating along the fit nematic director field of that reconstructed spindle from the minus end’s position to the pole (Figure 6A). We then calculated the distance *s* between the KMT’s minus end position and the pole along this streamline, with *s* = 0 for minus ends at the pole. A density histogram constructed by binning together all minus ends positions with respect to *s* reflect two distinct effects: 1) variations of KMT minus end positions along *s* within a k-fiber; 2) variations of the number of k-fibers along *s*. We wished to study the former, not the latter, so we focused on an alternative distribution: the density distribution of KMT minus ends along streamlines whose plus ends were upstream of that position. To construct that distribution, we first calculated the density of minus ends in a small bin within 0.1µm of the pole along the streamline trajectories. To find the density of the minus ends in the next 0.1µm bin upstream, we multiplied the KMT density in the first bin by the ratio of the number of KMT minus ends in the second bin with plus ends more than 500nm upstream from the second bin to the number of KMT minus ends in the first bin with plus ends more than 500nm upstream from the second bin. We then iterated this procedure along the streamline trajectory to produce the density distribution of KMT minus end along streamlines whose plus ends were upstream of that position (Figure 6B).

### Deriving mass conservation equation for KMT minus ends to calculate the KMT minus end speed *v*(*s*)

We performed a mass-conservation flux analysis on the KMT minus end density distribution (measured from the EM reconstructions) to predict the speed of the KMT minus ends throughout the spindle (Figure 6C). We assumed that the KMTs in metaphase are in steady state and move along streamlines, which implies that the fluxes associated with KMT gain, motion and loss must balance at every position along the streamlines. We considered a region along a streamline between positions *s* and *s* + *ds*, and defined the fluxes associated with KMT minus end gain, motion and loss in this region as:

1. **Gain**: New KMTs join the fiber with their minus ends at position *s* along streamlines at rate *j*(*s*). The form of *j*(*s*) depends on the choice of a model for how KMTs are recruited to the kinetochore and is discussed in more detail below.
2. **Motion**: KMT minus ends move into the region with flux *v*(*s* + *ds*) *n*(*s* + *ds*) and move out of the region with flux *v*(*s*) *n*(*s*), where *n*(*s*) is the density of KMT minus ends at position *s*, and *v*(*s*) is the speed of KMT minus ends at position *s*. Subtracting these terms and taking the limit *ds* → 0 gives the motion flux as 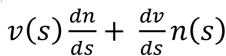
3. **Loss**: KMTs detach from the kinetochore and depolymerize at rate *r*. Our photoconversion experiments revealed that the lifetime of KMTs was independent of their position in the spindle bulk (Figure 3G), so we took *r*. to be constant (i.e. independent of *s*). We set *r* to be the inverse of the average lifetime of KMTs in the spindle bulk measured in the photoconversion experiments: i.e., *r* = 0.4 min^-1^

Since the KMT minus ends are in steady state, these three fluxes must sum to zero everywhere. This gives us a steady state mass conservation equation:

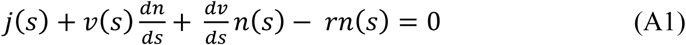

### Defining the *j*(*s*) gain flux term

The form of the *j*(*s*) gain flux term depends on the KMT recruitment model (Figure 6D). If all KMTs result from de novo nucleation at kinetochores (i.e., the nucleate at kinetochore model), then, by assumption, *j*(*s*) = 0 at all locations in the spindle bulk. Alternatively, if KMTs result from non-KMTs that bind the kinetochore (i.e., the capture from spindle model), then *j*(*s*) ≠ 0. For a non-KMT to bind a kinetochore, it must first be nucleated and then grow far enough to contact a kinetochore. Non-KMTs turnover in ∼0.25 min and move at a speed of ∼1 μm/min (Figure 3), so we estimate that they travel only ∼0.25 μm before depolymerizing. Thus, since non-KMTs are not expected to significantly move over their lifetime, we take the inferred density of non-KMT minus ends along streamlines, *n_NK_* (*s*) (Figure 6s2), as an estimate of the non-KMT nucleation rate along streamlines. The length distribution of non-KMTs is observed to be exponential, with a mean length of *l_NK_* = 1.9 ± 0.1 *μm* (Figure A2). Thus, if a non-KMT nucleates at position *s* along a streamline, the probabilities that it grows far enough to reach a kinetochore located at position *s*_0_ is proportional to 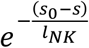. Taken together, this leads to 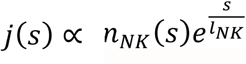 for the capture from spindle model, where the dependence on the position of the kinetochore is absorbed into the constant of proportionality.

### Integrating the mass conservation equation (A1) to find minus end speed predictions

We set a no-flux boundary at the pole to integrate the mass conservation equation (A1). The no-flux condition at the pole implies that either *n*(0) = 0 or *v*(0) = 0, reducing the mass conservation equation to:

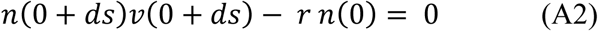

*n*(0) = 0 therefore requires that *n*(0 + *ds*) *v*(0 + *ds*) = 0 which reproduces the no-flux boundary condition at the position *ds*. Iterating this procedure produces a trivial solution that *n*(*s*) = 0 everywhere. Since we observed a non-zero minus end distribution, we used the *v*(0) = 0 condition instead. Using this *v*(0) = 0 condition we integrated equation (1) numerically to find the KMT minus end speed predictions from the nucleate at kinetochore and capture from spindle recruitment models (Figure 6E).

### Simulated 2D confocal imaging of a photoconverted line

We simulated the motion of tubulin after photoconversion in both KMTs and non-KMTs and the reincorporation of depolymerized tubulin in the simulation spindle for each of the three reconstructed cells (Figure A1). We assumed that the dynamics were the same along all streamlines and simulated the motion of photoconverted tubulin in KMTs and in non-KMTs along a streamline. We calculated the tubulin profile along the spindle axis from each streamline and then combined the results from the different streamlines. We finally added a background profile from depolymerized photoconverted tubulin that reincorporated throughout the spindle to produce a final line profile for analysis.

#### KMTs

We simulated the gain, motion, and loss of individual KMTs along streamlines at discrete timesteps. At each simulation timestep, we generated newly recruited KMTs with Poisson statistics. The KMT plus end positions were selected from the distribution of kinetochores along streamlines (Fig 7S1). For kinetochore nucleated KMTs, the minus ends started at the same location as the plus end. For spindle captured KMTs, the minus ends position was drawn from the probability that a microtubule would nucleate times the probably it would reach the kinetochore 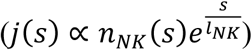. Newly encorporated tubulin polymerized at the KMT plus ends while the minus ends move backwards along the streamline towards the pole with an experimentally inferred speed *v*(*s*). In the spindle bulk, the minus ends moved at the same speed that the tubulin incorporated at the plus end. When KMT minus ends entered the pole region, at *s_p_* = 1.5*μm* upstream from the pole, tubulin continued to polymerize at the same rate as at the boundary, but the minus ends began to depolymerize at speed *v_tread_*(*s*) = [*v*(*s_p_*) − *v*(*s*)]θ(*s_p_* − *s*), where θ(*s*) is the Heavyside step function. The tubulin in a KMT therefore moved at speed *v_tub_*(*s*) = *v*(*s*) + *v_tread_*(*s*) while the minus ends moved at the experimentally inferred speed *v*(*s*). After an exponential drawn lifetime with mean 1/*r* = 1/0.4 min = 2.5 min, the KMTs detach from the kinetochore and are removed from the simulation.

To simulate the motion of photoconverted tubulin, we calculated the intensity of the photoconverted tubulin along streamlines with a modified Cauchy profile 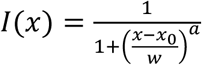.

Based on fits to the experimental line profile immediately after photoconversion, we set *a* = 1.7 and *w* = 400nm (Figure 7s2). The KMTs were pre-equilibrated for 20 minutes of simulation time before the simulation line was drawn to ensure the KMTs were in steady state. We projected the simulated photoconverted tubulin intensity along the spindle axis and summed the contribution of each KMTs along each of the spindle streamlines to produce a KMT line profile.

#### Non-KMTs

For the non-KMTs, we calculated the initial intensity of photoconverted tubulin along a streamline by multiplying the density of non-KMTs along the streamline by a Cauchy intensity profile along the spindle axis. We then translated the entire profile along the streamline towards the pole at a uniform speed equal to the speed of KMT minus ends where the line was drawn *v*(*s*). The profile height decayed at a rate *r_NK_* = 4min^-1^ measured in the photoconversion experiment (Figure 3H). Changing the simulated speed of the non-KMTs did not significantly impact the measured speed of the KMTs after the final analysis (Figure A3). Like the KMTs, we simulated the motion of the peak along each streamline, projected onto the spindle axis and then summed the streamlines together to produce a line profile. We added the KMT and non-KMT profile together, normalizing the profiles so that the KMT to non-KMT intensity ratio was 4:1.

#### Reincorporated Background

Finally, we included the contribution of reincorporated tubulin from photomarked microtubules that depolymerized. We modeled the background as a constant tubulin profile whose height exponentially approached a plateau value 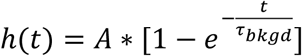 We determined the profile of reincorporated tubulin background from the average profile of tubulin along the spindle axis in cells with an mCherry:alpha-tubulin marker. (Figure A4). The height and timescale of the background profile were found using the photoconverted tubulin signal at the opposite pole in the photoconversion experiments. We fit a Gaussian to the photoconverted tubulin profile at the opposite pole. We then fit the height of the peak over time to determine the height and timescale of the background profile (Figure A5). The background incorporation took τ_*bkgd*_ = 80*s* and leveled off to *A* = 3% of the height of the original peak.

#### Fitting the motion and decay of the simulated peak

We summed the contribution of the KMTs, non-KMTs and background together and then convolved the line profile with a Gaussian with width 250nm to simulate the microscope point spread function. We then processed our simulated curves through the same algorithm we used to fit the experimental curves (see Methods): fit the pixels near the top of the peak to a Gaussian, fit the center of the Gaussian to a line to determine the velocity, fit the height of the Gaussian corrected for background to a dual-exponential to determine KMT and non-KMT stability.

#### Error Analysis

We repeated the simulations for each of the three EM-reconstructions. We used the measured KMT minus end distribution and spindle geometry from each individual spindle. We took the mean of the predictions from the three cells to find the model predicted speed of the photoconverted line (Figure 7B). We then took the standard error of the mean for the speed predictions from all three spindles to find the error in our model predictions.

### 3D Spindle Simulations

We simulated the gain, motion, and loss of discrete KMTs in each of the three reconstructed cells in 3D. At each timestep, we nucleated new KMTs at kinetochores by placing both the plus and the minus end at the same position within 200nm of the position of a kinetochore in the reconstruction. We then moved the minus ends of the existing KMTs towards the pole at the experimentally inferred speed *v*(*s*) along nematic streamlines in 3D. The nematic streamline for each KMT were found by calculating the 2D nematic streamline from the plus end position in the spindle-radial axis plane and rotating the spindle-radial axis plane about the spindle axis to the kinetochore position. This procedure produced a 3D streamline that was flat in the theta direction. When the KMT minus ends cross the pole boundary at *s_p_* = 1.5*μm* from the pole along a streamline, the minus ends begin to depolymerize causing tubulin to treadmill through the spindle at a speed *v_tread_*(*s*) = [*v*(*s_p_*) − *v*(*s*)]θ(*s_p_* − *s*), as in the 2D case. The KMTs detach from the kinetochore at a rate *r* = 0.4min^-1^ and are removed from the simulation.

We compared the predicted lengths, orientations, and dynamics of the simulated and experimentally measured KMTs. We measured the lengths of the simulated KMTs from the distance between the plus and the minus end along the streamline trajectory. To compare the orientations of the simulated and reconstructed KMTs, we divided each KMT into short 100nm subsections and projected the subsections onto the spindle axis. We compared the fraction of the 100nm subsection lengths along the spindle axis in the simulation and experiment. We drew a plane of photoactivation tubulin perpendicular to the spindle axis with a Cauchy profile. We projected the tubulin intensity in a thin 1μm confocal z-slice onto the spindle axis to produce a line profile. The center position, width, and exponent of the profile were fit to a sample photoconverted line profile at t=0 min. We then tracked the converted tubulin in the spindle over 60s of simulated time and reprojected the confocal slice onto the spindle axis to compare the line profile with experimental converted line profile at t=60s.

**Figure A1:**
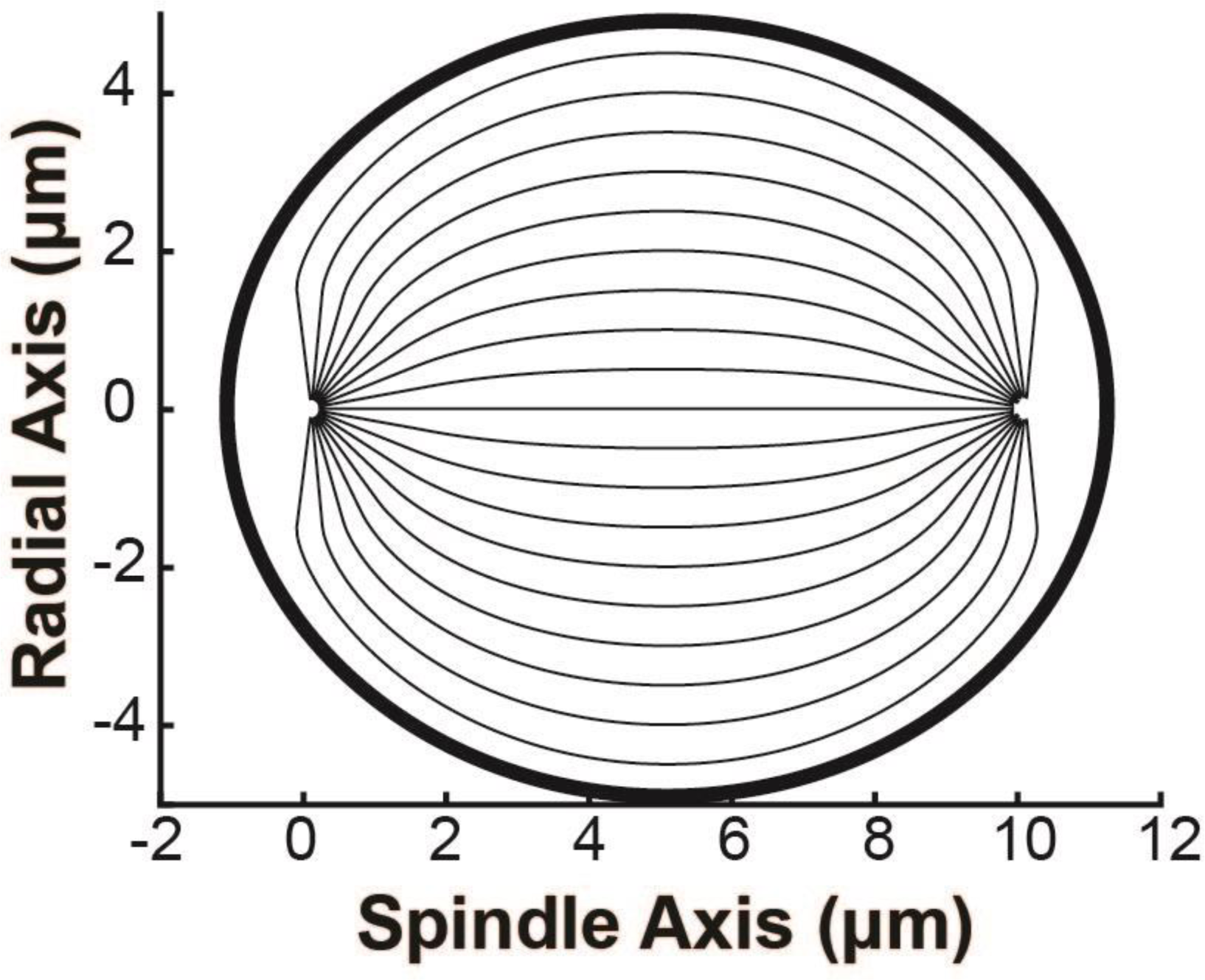
Sample geometry of spindle streamlines used in the simulation. Geometry of the spindle streamlines used in the simulations. The thin lines show the trajectories of nematic streamlines in the spindle bulk. The thick black line shows the elliptical boundary of the spindle.

**Figure A2:**
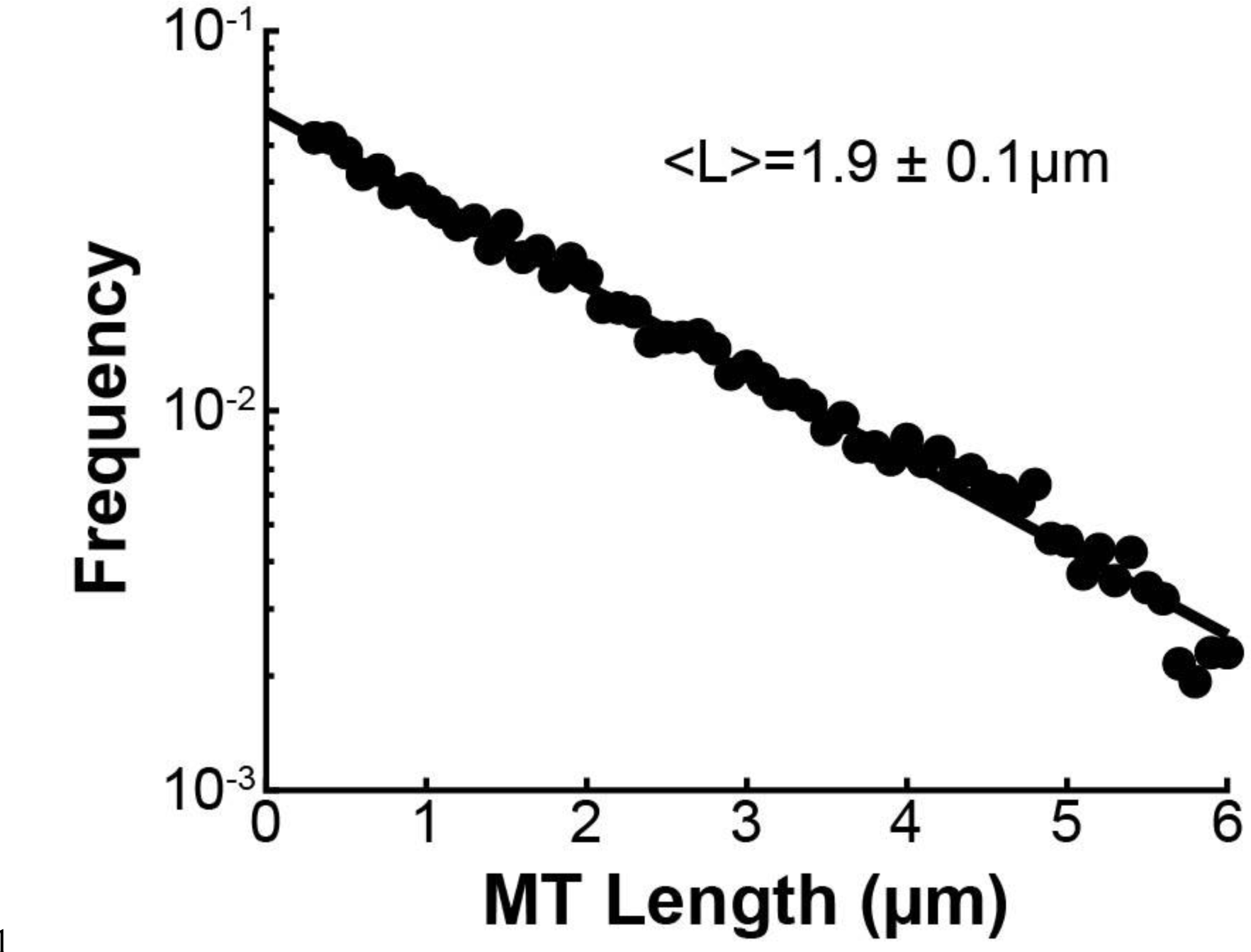
Length distribution of non-KMTs in the spindle. Binned histogram of the lengths of non-KMTs in three reconstructed mitotic HeLa spindle. Black dots: electron microscopy data; black line: exponential fit. Mean MT length is 1.9±0.1µm.

**Figure A3:**
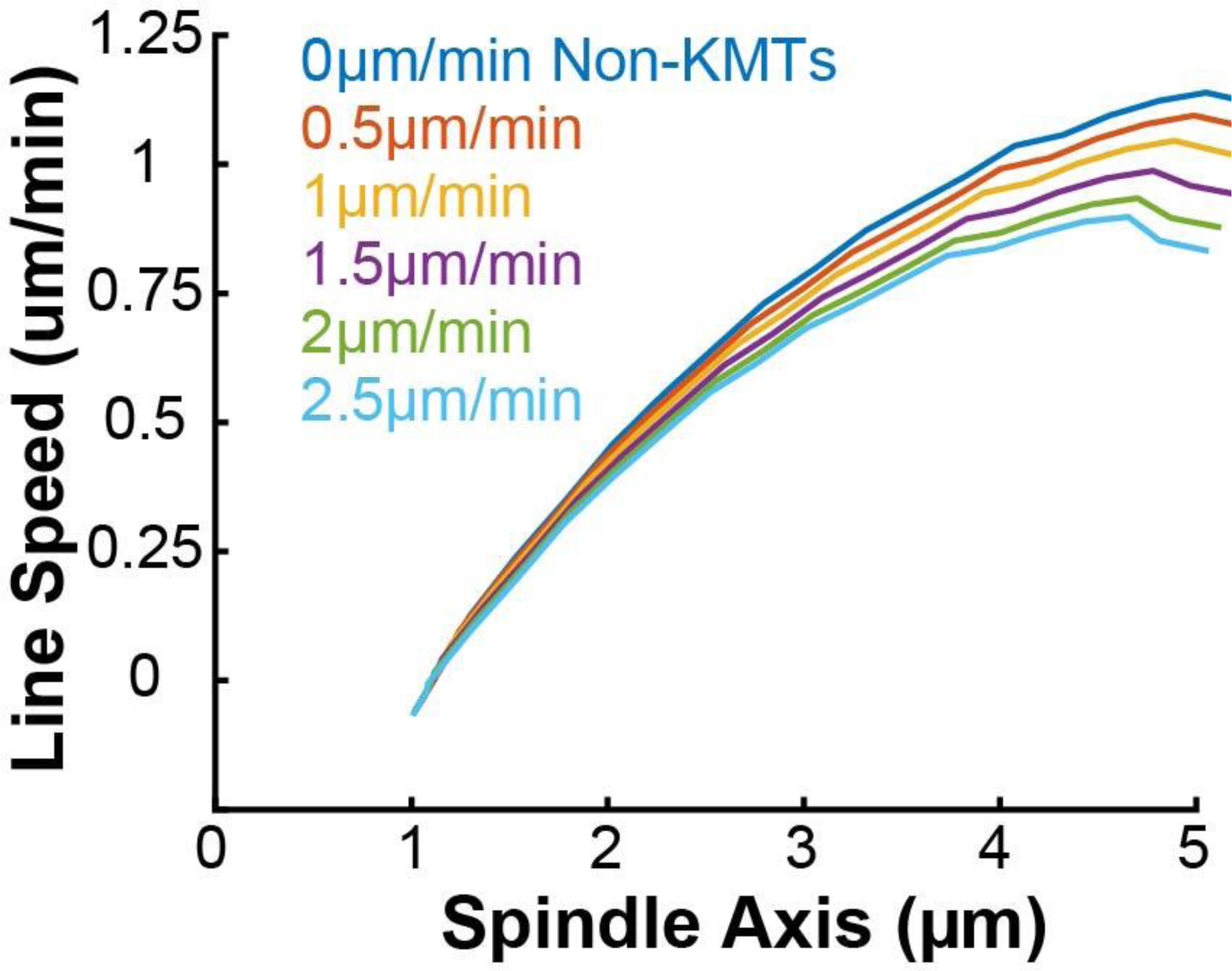
Predicted photoconverted line speed for various uniform non-KMT motion speeds. The speed of the non-KMTs was varied (assorted colors) in 0.5µm/min increments in a 2D confocal imaging spindle simulation.

**Figure A4:**
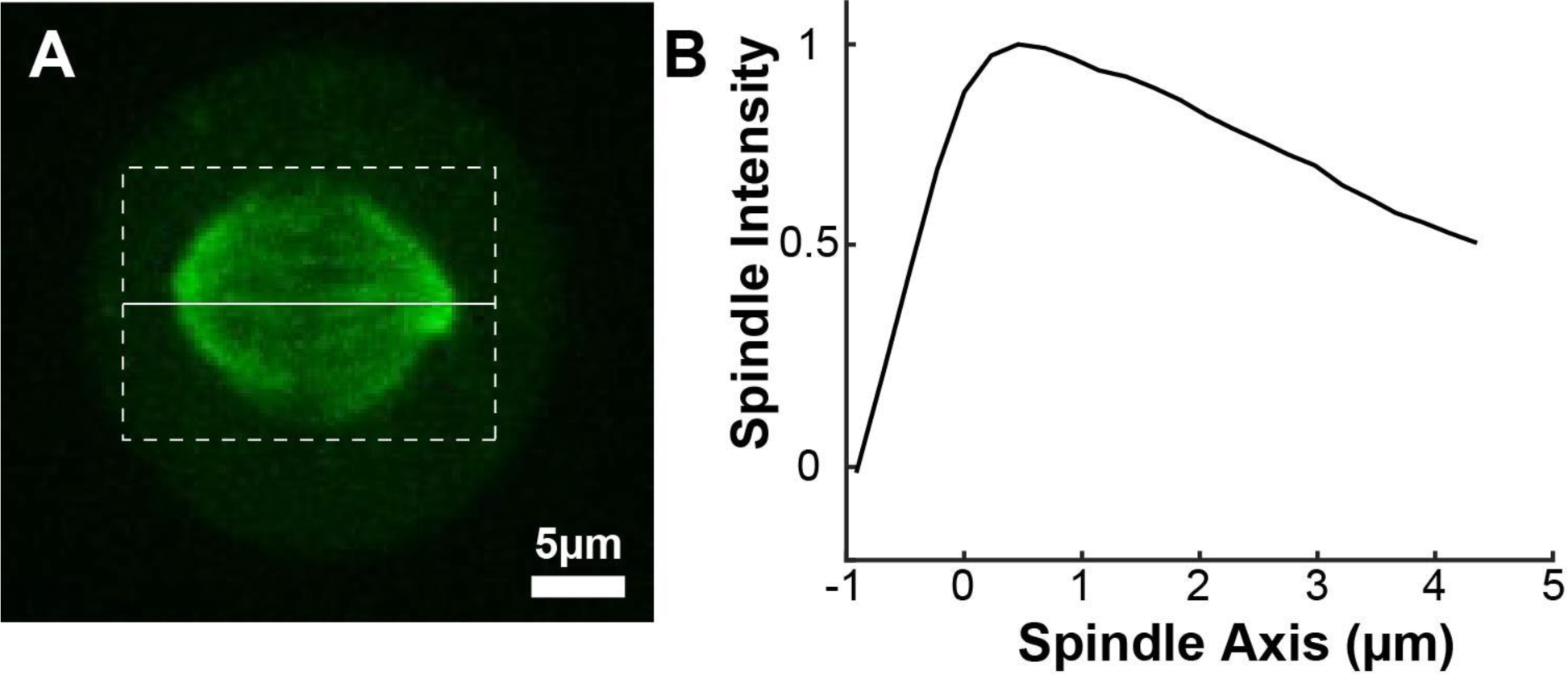
Spindle background profile. A) Sample representative spindle image (Green: mCherry:tubulin). B) The intensity of the tubulin marker projected onto the spindle axis and then averaged for n=72 half spindles. The spindle axis x=0 is located at the pole.

**Figure A5:**
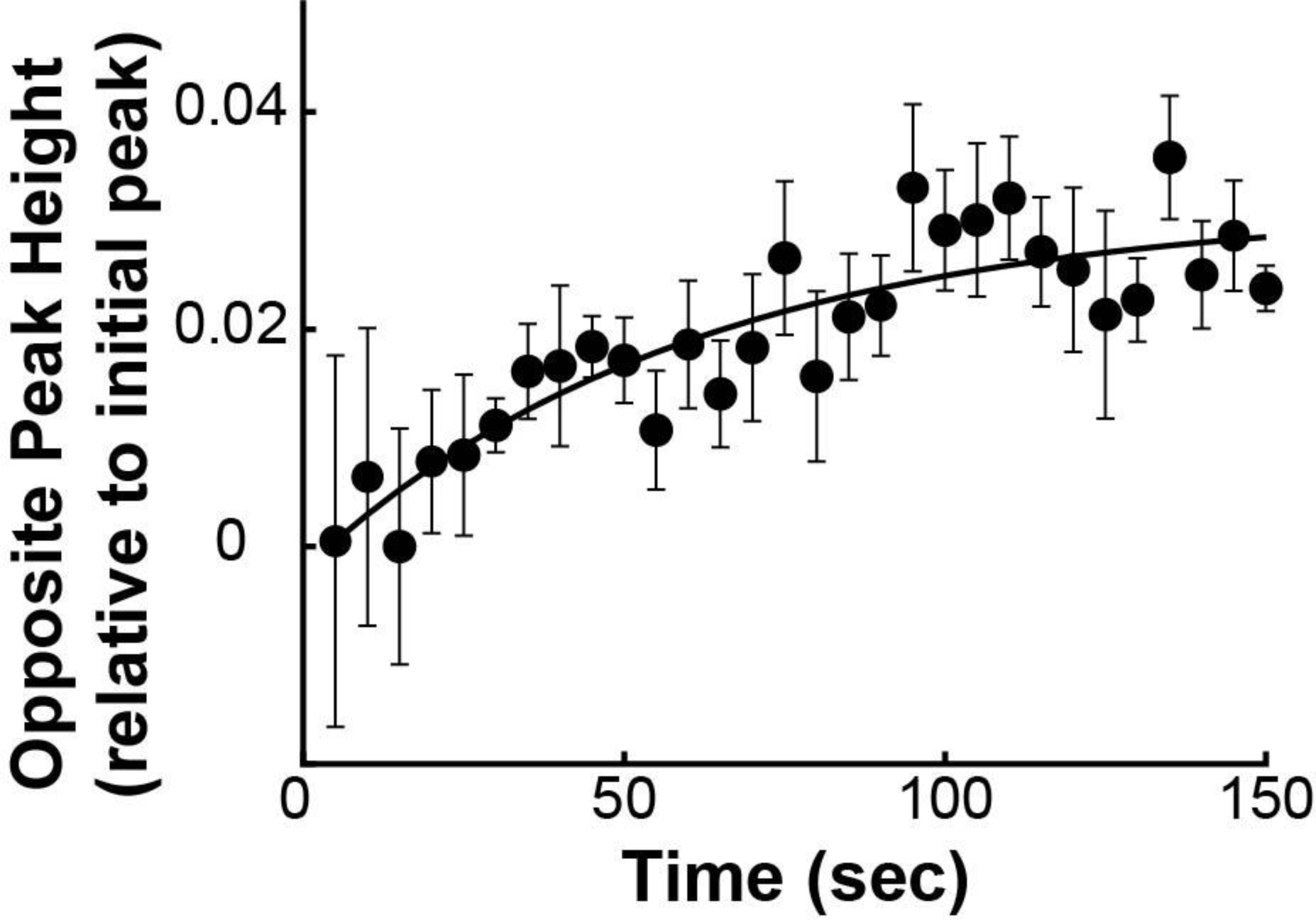
Height of the opposite pole over time. The peak height averaged from n=5 spindles displaying a clear opposite peak (black dots) is fit to an exponential (black line).

**Table A6:**
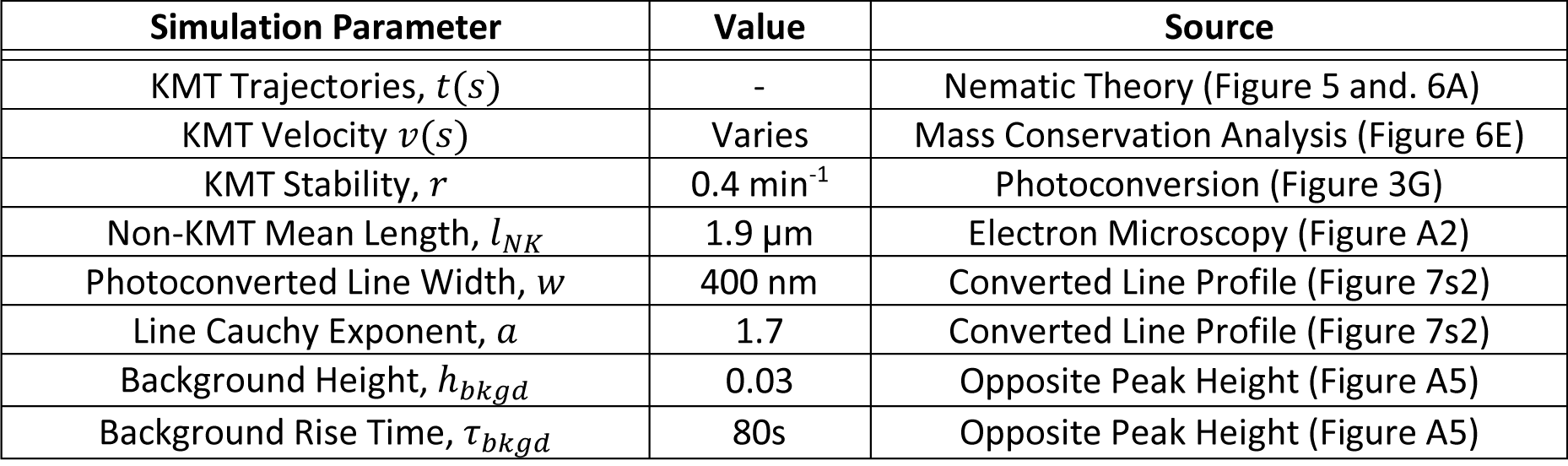
Parameters values and sources.

