## Supplementary figures and images for "Self-organization of kinetochore-fibers in human mitotic spindles"

### Video 8s1 - Simulated tubulin photoconversion in a 3D model spindle

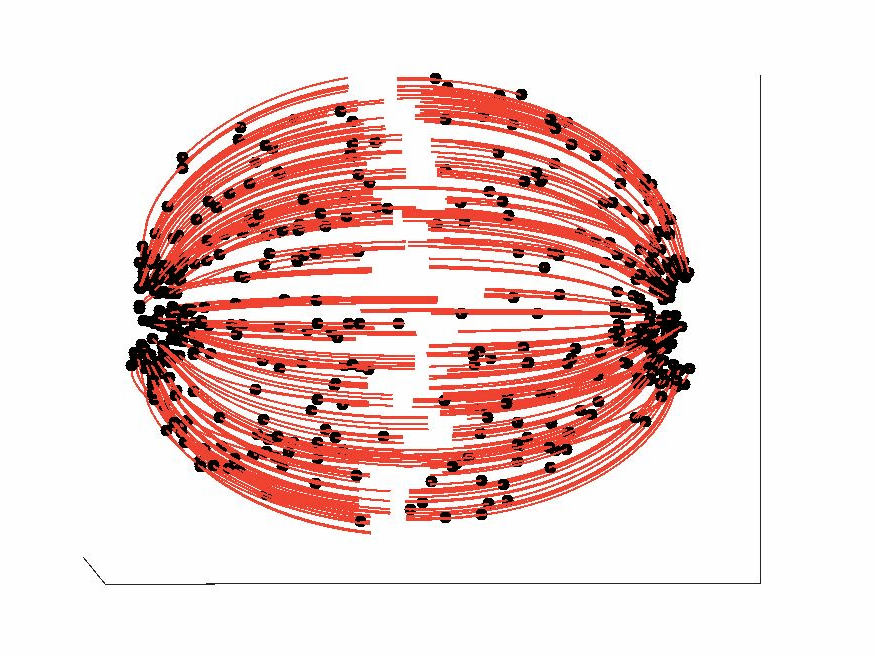
